# Genetically interacting mutations and mechanical stress affect the stochasticity of developmental eye defects in *yap1* mutants

**DOI:** 10.64898/2026.07.11.737910

**Authors:** Niccolo Fioritti, Mahum Shaikh, Giulia Cazzagon, Gareth Powell, Kate Turner, Lisa Tucker, Anna Mallucci, Evdokia Hadjieconomou, Stephen Carter, Daniel Wehner, Gilbert Weidinger, Richard J. Poole, Florencia Cavodeassi, Leonardo E. Valdivia, Rodrigo Young, Masazumi Tada, Alex Nechiporuk, Stephen W. Wilson, Gaia Gestri

## Abstract

Congenital abnormalities of eye formation show remarkably variable penetrance with phenotypes even varying between left and right eyes. Here we explore this phenomenon through analysis of the mechanistic basis of low penetrance retinal coloboma in zebrafish *yap1^nl13/nl13^* mutants and identification of factors that modify the probability of this phenotype. We find that the low penetrance stochastically occurring coloboma in *yap1^nl13/nl13^* mutants is due to rupture in the ventral retina at the point of apposition of the lips of the closing choroid fissure and that provision of wild-type Yap in the retinal pigment epithelium suppressed this phenotype. Decreasing actomyosin contractility increased the penetrance of coloboma whereas increasing myosin phosphorylation rescued the phenotype suggesting that altered mechanical properties of the RPE sensitize the eye to stochastic failure of choroid fissure closure. Genetic interaction screening revealed enhanced and synthetic eye phenotypes in *yap1^nl13/nl13^*mutants upon abrogation of function of genes encoding extracellular matrix and other genes implicated in eye formation. Our data reveal that the variable penetrance of congenital eye abnormalities can be due to genetic and environmental factors impacting the stochastic variability inherent in the developmental processes underlying eye morphogenesis.

## Introduction

Although hereditary, congenital abnormalities of eye formation are a major cause of blindness in children, the mechanistic bases of such phenotypes remain poorly understood (ALSomiry, 2019; Harding and Moosajee, 2019; Plaisancié et al., 2019; Williamson et al., 2014). We know even less about the highly variable penetrance and expressivity of such phenotypes. Remarkably, it is even common for left and right eyes to express very different phenotypes despite carrying the same genetic lesions. The process of eye formation is, however, highly conserved across vertebrates and progress is being made in identifying genes, that when disrupted, lead to anophthalmia loss of eyes (loss of eyes), microphthalmia (small eyes) and coloboma (failure in closure of the optic cup) in both humans and animal models (Cardozo et al., 2023; Yoon et al., 2020; Zuber et al., 2003).

The primordium of the eyes, the eyefield, is a single domain of cells in the anterior neural plate. Once specified, cells destined to form left and right eyes evaginate laterally, splitting the eyefield and giving rise to the optic vesicles. These structures undergo further morphogenetic rearrangements to form optic cups with inner neural retina and outer retinal pigment epithelium (Cardozo et al., 2023; Casey et al., 2023; Cavodeassi, 2018; Fuhrmann, 2010; Ivanovitch et al., 2013). The optic cup has a transient fissure on the ventral side allowing ingress of blood vessels and exit of retinal axons; this choroid, or optic, fissure subsequently fuses to close the globe of the eye. For these events to occur successfully, complex, coordinated interactions between tissues of different developmental origins are required. For instance, coloboma can arise because of disrupted gene function in the neural retina or in adjacent periocular mesenchyme (ALSomiry, 2019; Bassett et al., 2010; Bryan et al., 2020; Gestri et al., 2018; Lupo et al., 2011).

The forces, cell movements and cell morphology changes responsible for shaping the vertebrate eye are beginning to be resolved. The nascent optic vesicle is populated by multi-potent progenitors that have the potential to acquire cell identities with very divergent shapes and behaviours. The prospective neural retina is composed of highly proliferative, columnar pseudostratified neuro-epithelial cells whereas cells of the prospective RPE flatten and stretch around the neural retina in a process that requires little cell proliferation (Moreno-Mármol et al., 2021). This flattening of the RPE is coordinated with the invagination of the optic vesicle to form the bi-layered optic cup with the outer RPE enwrapping the inner neuroretina. Atomic force microscopy has shown that the high levels of phosphorylated myosin in the RPE make this tissue more rigid than the neuroretina and it is believed that these differences in tissue stiffness are required for the correct invagination of the optic cup (Carpenter et al., 2015; Eiraku et al., 2012, 2011). It is proposed that the rigid shell of the RPE constrains the consequences of proliferation in the softer neuroretina leading to inward folding and cup formation (Moreno-Marmol et al., 2018).

As a central coordinator of mechanical feedback, the Hippo/Yap signalling translates tissue level tension into transcriptional outputs that govern both morphogenetic processes and cell fate determination and has conserved roles in various aspects of eye formation across species (Asaoka et al., 2014; Cabochette et al., 2015; Lian et al., 2010; J. B. Miesfeld et al., 2015; Silvano et al., 2025; Williamson et al., 2014). For instance, mutations in *yap* in zebrafish and mice have been associated with defects in the specification and maintenance of RPE; *yap1^mw48^* zebrafish mutants show loss of RPE cells, a phenotype aggravated by deletion of the *taz* gene and in mouse, loss of Yap function leads to a more extreme phenotype with RPE cells transdifferentiating into neural retina (Kim et al., 2016; J. B. Miesfeld et al., 2015; Neelathi et al., 2026). In humans, mutation in the Yap/Taz binding domain of TEAD 1 have been associated with Sveinsson chorioretinal atrophy with loss of photoreceptors and RPE around the optic nerve head, while heterozygous loss-of-function mutations in YAP cause incomplete penetrant coloboma and microphthalmia (DeYoung et al., 2022; Fossdal et al., 2004; Holt et al., 2017; Williamson et al., 2014). Coloboma phenotypes have not been described in mice *yap* mutants but do occur in *yap1*^nl13/nl13^ zebrafish mutants (Kim et al., 2016; Masson et al., 2020; J. B. Miesfeld et al., 2015). One remarkable feature of many of these Yap-related phenotypes is the variability between left and right eyes.

Differences in mutant allele combinations undoubtedly explains some of the variation in eye development phenotypes between individuals but does not, for instance, readily explain why left and right eyes in the same individual can show different phenotypes. To understand such phenotypic variability, it is necessary to resolve why the phenotypes arise and why the implicated developmental processes sometimes succeed and sometimes fail. To this end, in this study we performed high resolution 4D imaging to analyse the key cellular events that underlie the variably penetrant choroid fissure defects in *yap1^nl13/nl13^*mutants and explored the environmental and genetic factors that affected successful fissure closure.

We show that coloboma in *yap1^nl13/nl13^* mutants is due to a stochastically occurring catastrophic structural failure in the ventral retina at the point of apposition of the closing lips of the choroid fissure. Our discovery that restoration of Yap function in the RPE rescues choroid fissure closure demonstrates that the coloboma phenotype is likely due to reduced rigidity of the RPE shell around the forming neural retina. We find that the probability of the stochastic failure in choroid fissure closure is enhanced by environmental temperature, by genetic and pharmacological manipulations that alter phosphorylated myosin / tissue tension and by loss of function of genes encoding extracellular matrix proteins that invest the forming eye. Our results show that detrimental genetic and environmental factors can impact the inherent stochastic variability of specific morphogenetic events during eye formation leading to high variability in penetrance of developmental retinal defects.

## Methods

### Zebrafish lines and husbandry

*AB*, *TU*, *WIK* and *EKWILL* wild-type, *Tg(UAS:GFP-YAP1)^u313^* (this study); *Tg{rx3::Gal4-VP16}^vu271^*(Weiss et al., 2012)*; Tg(tfec:GAL4)* (J. B. Miesfeld et al., 2015); E1-bhlhe40:GFP (Moreno-Mármol et al., 2021); *Tg(UAS:mCherry)^mw60^* (Miesfeld and Link, 2014); *Tg(β-actin:HRAS-RFP)^vu119^* (Cooper et al., 2005); *yap1^nl13^* (J. B. Miesfeld et al., 2015); *Tg(vsx2:GFP)^nns1^*(Kimura et al., 2006); *lrp5^ulm21^* (this study); *lrp6^u348^* (Powell et al., 2024); *fz5^sgu1^* (Monfries et al., 2025); *apc^CA50a^ (*Paridaen et al., 2009); *mab21l2^u517^*(Wycliffe et al., 2021); *lama1^nl14^* (this study) zebrafish (*Danio rerio)* lines were bred and maintained according to standard procedures (Aleström et al., 2020; Westerfield, M., 1993). Wildtype and mutant embryos were raised at 28°C if not otherwise stated and staged according to (Kimmel et al., 1995). The following alleles were genotyped by KASP assays (K Biosciences, *lama1^nl14^*, *apc^CA50a^*, *lrp6^u348^*; *lrp5^ulm21^*) or Tera-ARMS PCR (*fz5^sgu1^, yap1^nl13^*, *mab21l2*^u517^) see Table S2. To prevent pigment formation, 0.003% phenylthiourea (PTU, Sigma) was added to the fish water starting between 10 and 22hpf. All experiments conformed to European Community Directive guidelines and British (Animal Scientific Procedures Act 1986) legislation for the experimental use of animals.

The focus for this study is the analysis of the *yap1^nl13^*line. This *yap1* allele carries a mutation in a splice acceptor site that results in aberrant splicing and introduce an early stop codon in the transactivation domain for all the described isoforms (J. B. Miesfeld et al., 2015).

### Constructs and new lines

To assess the requirement of Yap1 in different cell types, we generated a line carrying a bidirectional *UAS: GFP-YAP1* transgene that drives GFP transcription in one direction and *yap1* in the other. *yap1* cDNA was cloned from zebrafish gastrula embryos then inserted along with GFP (GFP-Yap1) in the pBR-Tol2-UAS vector, as described (Kajita et al., 2014). Using this construct, the *Tg(UAS:GFP-YAP1)^u313^*line was generated and called *Tg(UAS:YAP1)* hereafter. Similarly the human Mypt1 construct obtained from the Kimelman lab (Weiser et al., 2009) was inserted along with mRFP in the pBR-Tol2-UAS vector to generate a dUAS:myr-RFP;*mypt1*. The *lrp5^ulm21^*allele was generated by CRISPR/Cas9 (IDT) using a guide RNA targeting exon 2 of *lrp5* (Table S2). The mutation is predicted to produce a truncated protein of 56 amino acids, compared with the full-length protein of 1,616 amino acids. The *lama1^nl14^* allele was identified through a forward genetic screen of ENU-induced mutations.

### RNAseq extraction and analysis

RNAseq was performed on 3dpf embryos raised at 28°C. To obtain embryos at the same developmental stage, *yap1^nl13/+^* heterozygotes were kept apart in breeding tanks and embryos collected 30 minutes after divider removal. Mutant and sibling embryos were sorted by phenotype at 3dpf. For RNAseq analysis, we performed 2 biological replicates and 1 experimental replicate. Total RNA was isolated from 45 embryos. RNA extraction and analysis were performed as described (Turner et al., 2019).

### *In situ* hybridization, immunolabelling and TUNEL staining

Embryos were fixed in 4% paraformaldehyde and standard methodology was followed for *in situ* hybridisation analysis (Thisse and Thisse, 2007); immunolabelling (Turner et al., 2019) and TUNEL (ApopTag Kit (Chemicon International); (Young et al., 2019). To improve TUNEL staining, we used 30-minute PK treatment rather than in situ/immunostaining. In embryos with increased apoptosis, a 0.5X dilution of the standard NBT/BCIP mix was used to slow colour development.

Primary antibodies were as follows: rabbit anti-P-Myosin Light-Chain 2 (dilution 1:100, Cell Signaling, Cat# 3671P), mouse Zonula Occludens 1 (dilution 1:200 Thermo Fisher Cat# 33910), chicken anti-GFP (Dilution 1:500 Abcam Cat# ab13970). Secondary antibodies were: Alexa Fluor 633 anti-mouse, 488 anti-rabbit, and 488 anti-chicken (all 1:1,000, Invitrogen). For cryosections, embryos were first protected by sequential incubation in 15% then 30% sucrose in phosphate-buffered saline supplemented with 0.5% Triton X-100 (PBST) for 15-30′, embedded in OCT, stored at −70°C, and sectioned at 18μm using a Leica cryostat. The *arr3a; opn1; tcap in situ* probe templates were generated directly by PCR (see Table S1), while the *prss1* plasmid was provided by (Mayer and Fishman, 2003).

### Temperature dependence and selection of *yap1^nl13/+^* carrier fish

*yap1^nl13/nl13^* embryos show variable eye phenotypes ranging from coloboma to overtly normal eyes with penetrance of these and other phenotypes varying between different pairs of fish. We generally selected carriers giving rise to healthy embryos with low penetrance phenotypes at permissive temperatures. Depending on the parent fish and experimental paradigms, embryos were raised at three different temperatures (28°C/30°C/32°C) to obtain different proportions/severities of *yap1^nl13/nl13^*coloboma phenotypes. In most clutches raised at 28°C, coloboma was rare and very mild if present; the proportion showing phenotypes increased at 30°C and coloboma was fully penetrant at 32°C. Some *yap1^nl13/nl13^*mutants raised at 32°C showed heart oedema and so 30°C degrees was usually selected as an optimal temperature to exacerbate eye phenotypes without affecting general development.

For the F0 genetic interaction screen, we selected *yap1^nl13/+^*heterozygotes that showed almost no phenotype in their progenies when raised at 28°C. For analysis of *yap1^nl13/nl13^* coloboma phenotypes, embryos were raised at either 30°C/32°C to obtain significant numbers of mutants with the phenotype. Rescue experiments were performed both at 28°C and 30°C with no differences in outcomes observed at the two temperatures (data is shown for experiments at 30°C). RNAseq experiments were carried out with embryos raised at 28°C to avoid general activation of stress related genes.

### Imaging and Data Processing

4D confocal imaging of embryos transgenic for both the RPE transgene [Tg(E1- *bhlhe40*:GFP) Bovolenta 2020.] and the cell membrane Tg(β-*actin*:HRAS- RFP)^vu119^(Cooper et al., 2005) or the nuclear mRNA H2A-RFP were anaesthetised in 0.2% tricaine methanesulfonate (MS222, Sigma) in fish water and mounted in 1% low melting agarose gel in embryo medium. A Leica TCS SP8 microscope with a 10x/0.30 NA HC or 25x/XXNA water-immersion lens was used (Leica). Z-stacks were acquired every 5–20 min for up to 14h. Timelapse movies were analyzed using Fiji (ImageJ) and Imaris software.

### Genotyping and sequencing

Genomic DNA was extracted from zebrafish embryos or fin clips and used as template in PCR reactions (Powell et al., 2024). Genotyping of the *yap1^nl13^* mutation was performed following PCR analysis of genomic DNA followed by BbvCI restriction digestion to assess for the presence of BbvCI restriction site, abolished by the *yap1^nl13^* mutation (see Table S2). For each digestion reaction, 5μL of the PCR product were added to 5μL of reaction mix (reaction mix: 1μL 10x Cutsmart Buffer (NEB, Cat# B7204S); 0.2μL BbvCI restriction enzyme (NEB, Cat# R0601L, 3.8μL water) and incubated at 37°C overnight. PCR was run on a 1% agarose GEL. Latterly, a Tetra-ARMS PCR assay was performed (Table S2). This method employs allele-specific inner primers and common outer primers to distinguish wild-type and mutant alleles in a single PCR reaction and was used for *yap1^nl13^*, *fz5^sgu1,^* and *mab21l2^u51^*(Table S2).

*lama1^nl14^*, *apc^CA50a^, lrp5^ulm21^* and *lrp6^u348^* were genotyped using KASP genotyping assays, a fluorescence-based method for SNP and indel discrimination (Table S2). KASP genotyping assays were either selected from the zebrafish SNP panel or custom-designed based on flanking sequence data and obtained from LGC Biosearch Technologies (formerly K Biosciences). Each reaction contained two allele-specific forward primers and a shared reverse primer, with allele detection based on FAM or HEX fluorescence signals.

### Statistics

Statistics and graphs were prepared using R (version 4.4.1; [R Core Team (2024). R: A Language and Environment for Statistical Computing. R Foundation for Statistical Computing, Vienna, Austria. https://www.R-project.org/]), ggplot2 package (version 3.5.1 [H. Wickham. ggplot2: Elegant Graphics for Data Analysis. Springer-Verlag New York, 2016.]), superb package (version 0.95.19; [Cousineau, D., Goulet, M.A., & Harding. B (2021) Summary plots with adjusted error bars: The superb framework with an implementation in R Advances in Methods and Practices in Psychological Science, doi: https://doi.org/10.1177/25152459211035109]) and RStudio (version 2024.12.0+467; Posit Software, PBC). The threshold for statistical significance was set at *p* < 0.05.

Two-tailed Welch two sample *t*-tests were used to compare differences in mean values. Two sample Kolmogorov-Smirnov tests with Monte Carlo *p* value simulation were used to test differences between distributions. Equality of proportions was tested using a Qʹ test and *post hoc* modified Marascuilo procedure with Benjamini-Hochberg correction for multiple testing [G. A. Michael, A significance test of interaction in 2 x K designs with proportions. TQMP 3, 1-7 (2007)]. Fisher’s Exact test was used in cases where only two samples were being compared.

The confidence interval for a mean value was calculated using a normal assumption. Confidence intervals for proportions were also calculated using a normal assumption if 0.1 < p^ < 0.9 and/or n > 20; otherwise by the Wilson count method.

### Pharmacological treatments

Para-nitro-blebbistatin (Cat# DRN111 Optopharma, Malnasi-Csizmadia 2014) was suspended in 100% DMSO to a final concentration of 5mM. Treated embryos were exposed to para-nitro-blebbistatin diluted from stock in E3 medium to the desired concentration. Titration experiments showed that 2μM was the highest subcritical concentration of Blebbistatin that led to lack of developmental defects in wild-types raised from 5hpf to 48hpf in continuous exposure to the drug.

### Morpholino, RNA and plasmid injections

0.25 ng of a splice-blocking morpholino against *mypt1*; 50pg of H2B:RFP mRNA (Anton et al., 2018) or 20pg of dUAS:myr-RFP;*mypt1* plasmid were injected at 1 cell stage in 1nl volumes.

### Generation of F0 crispants

Guide RNAs to target *shisa2a; shisa2b*; *axin1; apc; col9a2; col9a1b* were generated as described (Kroll et al., 2021). A protocol describing how to generate F0 crispant larvae is available at 10.17504/protocols.io.5qpvo52wdl4o/v3.Three synthetic RNA guides per gene were designed (Table S2). A subset of the injected embryos was genotyped by Illumina MiSeq data analysis (Table S2) to confirm the presence of gene editing (Kroll et al., 2024).

### Plastic re-use

Embryos were collected and grown in re-used petri dishes. To re-use petri dishes after exposure to fish water; 1%PTU or 1%Tricaine, they were soaked in hot water for at least 20 minutes (10.17504/protocols.io.yxmvmbxkng3p/v1).

To reduce plastic use, tips and pipettes for PBS/Ethanol/Methanol/fish water/SCC/PFA were used multiple times for different rounds of *in situ*, immunostaining and TUNEL (https://dx.doi.org/10.17504/protocols.io.81wgboxp3lpk/v1).

To minimize tip usage, washed gel-loading tips were reused as described (dx.doi.org/10.17504/protocols.io.e6nvw495dlmk/v1).

## Results

### Temperature stress during optic vesicle outgrowth sensitises *yap1^nl13/nl13^* mutant eyes to developing coloboma

The penetrance of coloboma in *yap1^nl13/nl13^* mutants is temperature sensitive; when clutches of embryos were raised at 28°C, coloboma was variably present in 10 to 60% of *yap1^nl13/nl13^* mutants and, when present, was usually unilateral (∼95%; Fig.1B,D). Penetrance was consistently higher at 30°C and when embryos were raised at a higher, stress-inducing, temperature (32°C; (Pype et al., 2015), penetrance of coloboma was 100% and bilateral (Note that full penetrance of coloboma in *yap1^nl13/nl13^*mutants would manifest as approximately 25% of embryos in a clutch showing the phenotype; Fig.1C,D).

**Fig 1:**
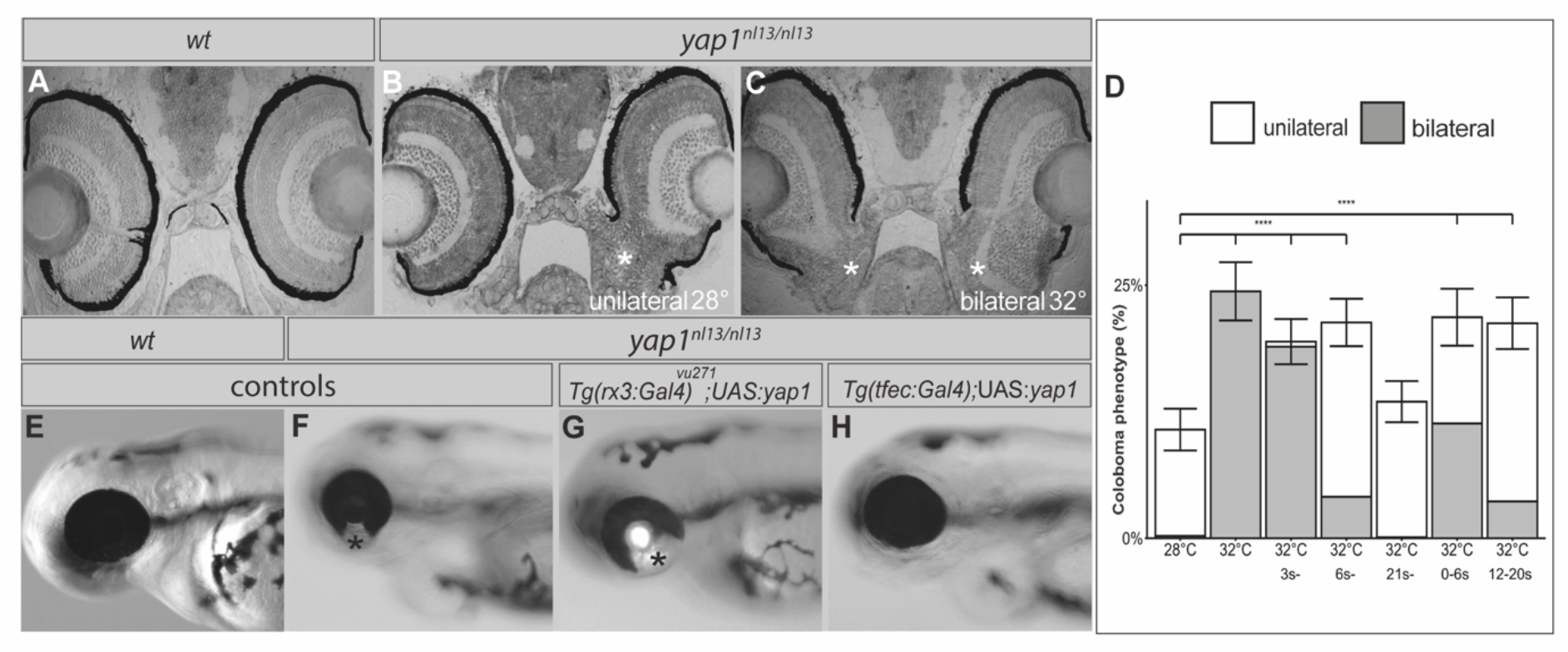
Penetrance of coloboma in *yap1^nl13/nl13^* mutants is sensitive to high temperature during early eye formation and is rescued by expressing wildtype Yap1 in the RPE. (A-C) 15 µm sections of *yap1^+/+^* wildtype (A) and colobomatous (asterisks) eyes in *yap1^nl13/nl13^* mutants raised at 28°C (B) and 32°C (C). D) Bar graph showing the percentage of embryos with coloboma in progeny of *yap1^nl13/+^* incrosses at different incubation temperatures and different stages (s = somite stage). Clear or grey-shading indicates unilateral or bilateral coloboma respectively. Equality of proportions Qʹ test (*χ*^2^=92.9, degrees of freedom = 6, p=7.54×10^-18^, n=867, 857, 1205, 829, 1173, 989 and 1090 embryos respectively) and *post hoc* modified Marascuilo procedure with Benjamini-Hochberg correction for multiple testing. ****p≤0.005; not all statistically significant comparisons shown for clarity. Error bars indicate the 95% confidence interval for the proportion of embryos with coloboma (unilateral and bilateral). (E-H) Lateral views of eyes of 3dpf fry: *yap1^+/+^*wildtype (E); *yap1^nl13/nl13^* mutant (F) (asterisk shows coloboma); *yap1^nl13/nl13^* mutant overexpressing wildtype *yap1* in the neural retina (G; *Tg(rx3:Gal4)^vu271^* driver) or in the RPE (H, *Tg*(*tfec*:*Gal4*) driver).

We used temperature shifts to assess at which developmental stages the penetrance of the coloboma phenotype was sensitive to temperature changes. Surprisingly, embryos from in-crosses of *yap1^nl13/+^* fish maintained at 32°C from stages when the optic cup is invaginating (21 somite stage (s); 21s) and throughout the period of fissure closure, did not show an increase in penetrance of coloboma (Fig.1D). Coloboma was only fully penetrant if embryos were raised at 32°C from the onset of optic vesicle evagination from the eye field (continuously from 3s or between 0-6s; Fig.1D), whereas penetrance progressively decreased when embryos were incubated at 32°C at later stages (12-20s Fig.1D). Consequently, temperature stress during outgrowth of the optic vesicle sensitises developing eyes to showing a fully penetrant coloboma at a later stage.

### Coloboma in *yap1^nl13/nl13^* mutants is suppressed if Yap1 function is restored in the RPE

Coloboma can arise from defects either in the neural retina or in the periocular mesenchyme (Gestri et al., 2018; Lupo et al., 2011; Matt et al., 2005) and *yap1* is expressed in both tissues as well as in the RPE (J. B. Miesfeld et al., 2015; Williamson et al., 2014). To determine in which cell type Yap1 function is primarily required to ensure closure of the choroid fissure, we restored wild-type *yap1* function to discrete cell populations in *yap1^nl13/nl13^* mutants.

Expressing exogenous wild-type Yap1 (together with GFP) throughout the forming optic vesicles using the *rx3: Gal4^vu271^* transgene failed to rescue (and indeed increased) coloboma in *yap1^nl13/nl13^*mutants raised at 30°C (Fig 1G; coloboma in n=32/32 GFP+ *yap1^nl13/nl13^*mutants; n=21/30 GFP- *yap1^nl13/nl13^* mutants). Wild-type siblings overexpressing *yap1* in the whole optic vesicles showed larger eyes and RPE defects (Fig.S1A-D). Coloboma was not rescued either when Yap1 was specifically expressed in the periocular mesenchyme using the *-5kblmx1b.1: Gal4-VP16^mw42^* transgene (data not shown; coloboma in n=14/19 *yap1^nl13/nl13^* GFP+ embryos; n=10/14GFP- *yap1^nl13/nl13^* mutants). Given that *yap1* is strongly expressed in the RPE and RNAseq analysis revealed robust gene expression changes in genes including *arr3* and *rpe65a* in the outer retina of *yap1^nl13/nl13^*mutants (Fig.S2, Table S1), we expressed wild-type *yap1* in the RPE using the *Tg(−2.7 kb tfec:Gal4)* driver (Lister et al., 2011; Miesfeld and Link, 2014). None of the GFP+ mutants obtained from *Tg (- 2.7kb tfec:Gal4);yap1^nl13/+^* x *Tg(UAS:yap1); yap1^nl13/+^* crosses showed coloboma (Fig 1H; n=0/36 GFP+ *yap1^nl13/nl13^* mutants; 18/26 GFP- *yap1^nl13/nl13^*mutants). Providing that wild-type *yap1* within the RPE is therefore sufficient to rescue choroid fissure closure in *yap1^nl13/nl13^* mutants.

### RPE specification and expansion are not overtly affected in *yap1^nl13/nl13^* mutants

As restoration of *yap1* function in the developing RPE rescued choroid fissure closure, we assessed if early events in RPE development such as RPE specification (from around 10s) and cell shape transition from a pseudostratified neuroepithelium to a squamous monolayer (from about 20s; (Moreno-Mármol et al., 2021) were affected in *yap1^nl13/nl13^* mutants.

Time-lapse studies showed that RPE specification and cell flattening occurred normally in *yap1^nl13/nl13^* mutants (Fig.2; Movie S1, S2 n=14 wt; n=6 *yap1^nl13/nl13^*). Emerging RPE cells were initially pseudostratified (Fig.2 A, B t=0) subsequently organising into a monolayer and acquiring cuboidal morphology (Fig.2 A,B t=40’). As optic vesicle invagination proceeded, RPE cells progressively flattened and acquired squamous morphology around the entire eye within a couple of hours (Fig.2A,B t=1.20, t=2h). These events were indistinguishable between wildtypes and *yap1^nl13/nl13^* mutants raised at 32°C. Similarly, we detected no significant changes in the numbers of RPE cells both at the time of RPE specification and optic cup formation or in the extent of coverage of the neural retina by the expanding RPE between wildtypes and *yap1^nl13/nl13^* mutants (Fig.2C-E n=12 wt; n=6 *yap1^nl13/nl13^*Movie S3, S4 and Fig.S3). These results show that the *yap1^nl13^*mutation does not overtly affect RPE specification, expansion, and cell shape remodelling during the early phases of RPE genesis.

**Fig. 2:**
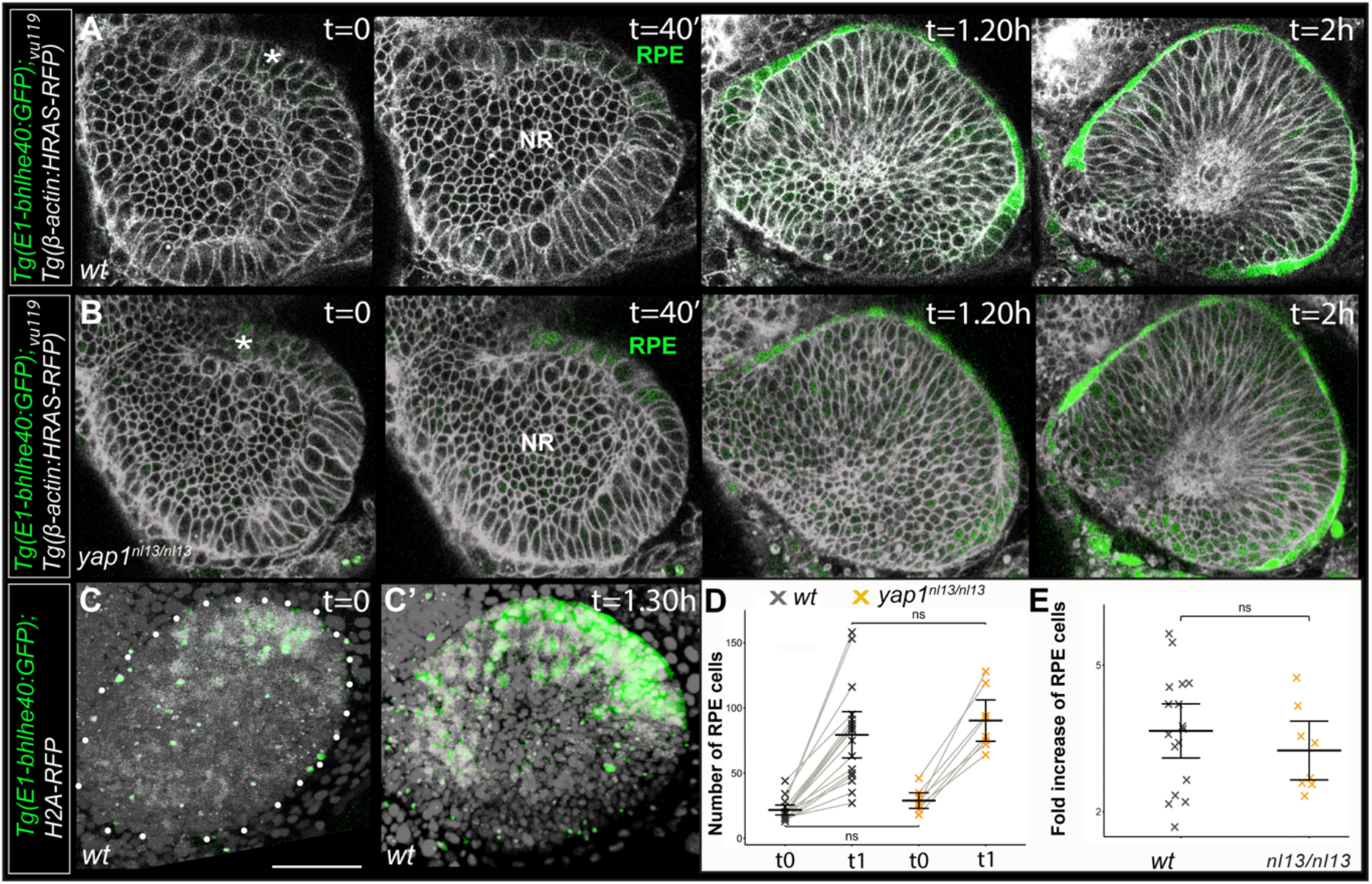
RPE specification and cell shape/cell number are not affected in *yap1^nl13/nl13^* mutants. A-B) Lateral views of developing optic vesicles in control (A) and *yap1^nl13/nl13^*embryos (B) at the time of RPE specification (t=0; 12s stage); cuboidal shape acquisition (t=40’); flattening of the RPE (t=1.20 and t=2h). RPE cells are marked by *E1-bhlhe40:GFP* expression (green) and cell membranes visualized in grey by *Tg(β-actin:HRAS-RFP)^vu119^* expression. C-C’) Lateral view of the optic vesicle at the time of RPE specification (C, t=0) and 1.5 hrs later (C’, t=1.30h). RPE cells express *E1-bhlhe40::*GFP(green) and all nuclei are stained by RFP-tagged H2A (pseudo-coloured in grey). Scale bar: 50 μM. D) Plots showing numbers of RPE nuclei at the stage of RPE specification. There is no significant difference between *yap1^nl13/nl13^* and control eyes (t0: Welch two sample, two-tailed *t*-test, *t* = -1.98, degrees of freedom = 12.80, p = 0.07; t1: Welch two sample, two-tailed *t*-test, *t* = -0.89, degrees of freedom = 21.04, p = 0.38). Crosses represent individual embryos, bars represent mean and 95% confidence intervals (cell count t0, mean ± 95% confidence interval: wildtype siblings = 21.7 ± 3.8, n = 17 embryos, 369 cells; *yap1^nl13/nl13^* = 28.9 ± 6.0, n = 8 embryos, 231 cells; cell count t1, mean ± 95% confidence interval: wildtype siblings = 79.4 ± 17.7, n = 17 embryos, 1349 cells; *yap1^nl13/nl13^* = 90.3 ± 15.9, n = 8 embryos, 722 cells;). ‘ns’ indicates not statistically significant. E) Plots of fold increase in RPE cell number calculated as the ratio between t0 and t0+t1. The fold increase values were not significantly different between wildtype and *yap1^nl13/nl13^* embryos (Fold increase, mean ± 95% confidence interval: wildtype siblings = 3.6 ± 0.5, n = 17 embryos; *yap1^nl13/nl13^* = 3.2 ± 0.6, n = 8 embryos; Welch two sample, two-tailed *t*-test, *t* = 0.96, degrees of freedom = 18.17, p = 0.35).

### Stochastically occurring collapse of the closing lips of the choroid fissure causes coloboma in *yap1^nl13/nl13^* mutant eyes

To elucidate the morphogenetic events that lead to coloboma in *yap1^nl13/nl13^* eyes, we performed high temporal and spatial high resolution confocal imaging of eyes expressing β-*actin*: HRAS-RFP^vu119^ and an RPE specific transgene, E1-*bhlhe40:*GFP, between 30-56hpf. The nasal and temporal lips of the choroid fissure approach each other from around 30hpf with subsequent fusion spreading bi-directionally in a zipper-like manner, distally along the ventral retina and proximally along the optic stalk (Fig.3A; (Gestri et al., 2018)). Up to 36/42hpf, *yap1^nl13/nl13^* eyes were indistinguishable from wild-type eyes (Fig.3B, C) with the RPE fully enveloping the invaginating optic cup. However, at the stage when the nasal and temporal lips of the choroid fissure became directly apposed in wild-type eyes, we observed rapid, ventrally directed rupture in some *yap1^nl13/nl13^* mutant eyes, leading to coloboma (n=4/7 imaged mutant eyes; Fig.3B, C; Movies S5-S7). In some cases, fragmentation of GFP fluorescence suggested RPE cells may be dying (Fig.3C, t=3h) though we did not observe obviously increased apoptosis in *yap1^nl13/nl13^* mutants at this stage using TUNEL staining (Fig.S4 B, E and data not shown). This paucity of identifiable apoptotic cells may be due to the presence of increased numbers of macrophages clearing dying cells in *yap1^nl13/nl13^*mutants (Fig.S2J/J’).

**Fig. 3:**
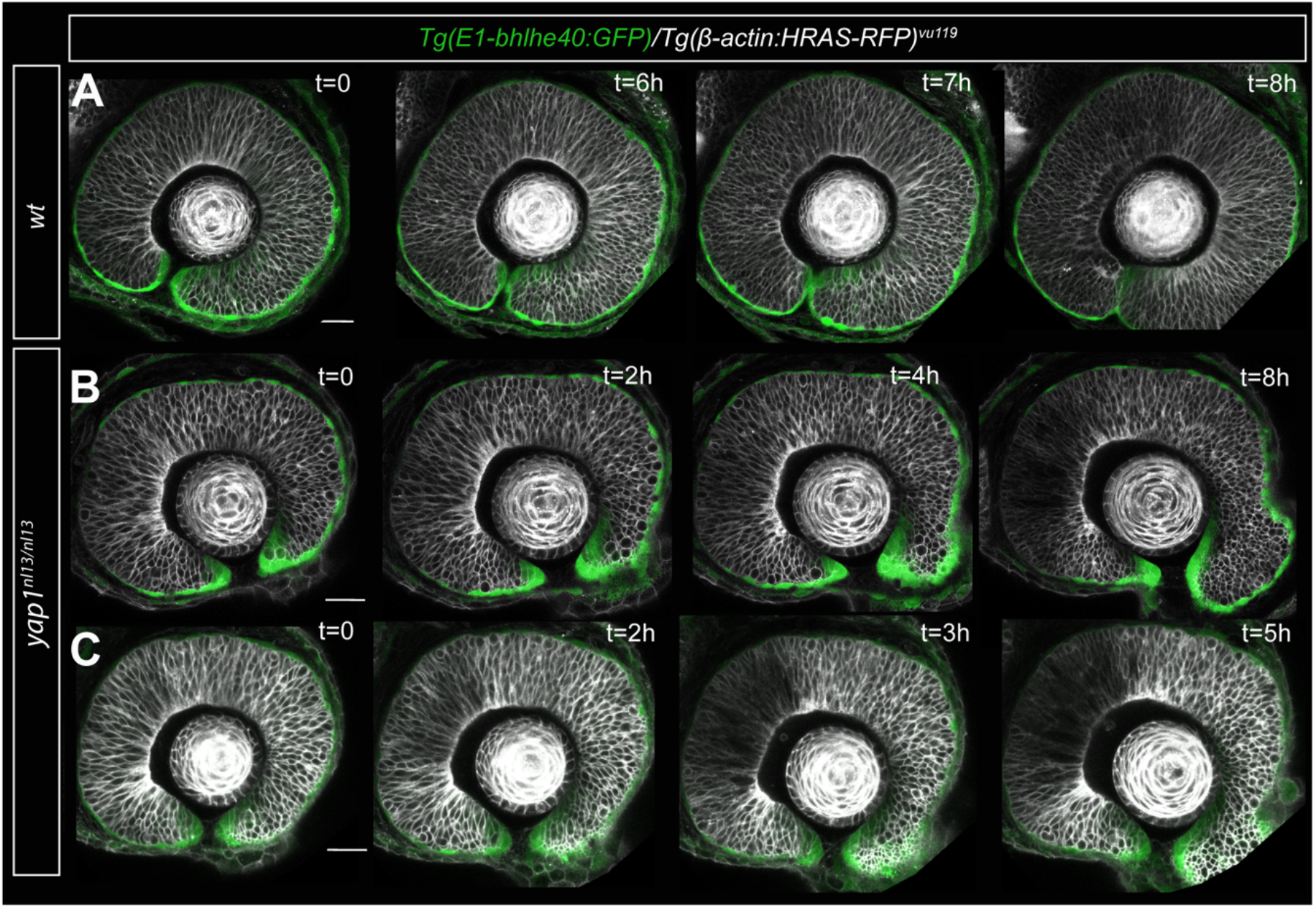
Choroid fissure collapse in *yap1^nl13/nl13^* mutant eyes. A-C) Tissue organization and behaviour during choroid fissure closure. Image stills from 16h (30- 56hpf) time-lapse movies (movies S4-S6) of wildtype (A) and *yap1^nl13/nl13^* mutant (B,C) eyes showing RPE cells expressing Tg(E1-*bhlhe40*:GFP) and cell morphologies (grey) visualised with Tg*(β-actin:HRAS-RFP)^vu119^*. Scale bar 30 µm.

### Compromising myosin phosphorylation leads to full penetrance coloboma in *yap1^nl13/nl13^* mutant eyes

The stiffness of the RPE increases during optic cup maturation and differential stiffness between RPE and neural retina supports invagination of optic cup organoids (Eiraku et al., 2012). Consequently, the coloboma phenotype observed in *yap1^nl13/nl13^*mutants could potentially be due to the inability of the *yap1^nl13/nl13^* RPE to generate a sufficiently stiff shell around the ventral NR. Phosphorylated myosin can increase tissue stiffness by enhancing the contractility of the cortical actomyosin complex (Munjal and Lecuit, 2014).To determine if we could observe altered actomyosin activity in the RPE of *yap1^nl13/nl13^* mutants, we visualised cortical myosin phosphorylation (pMLC) in RPE cells with ZO1-labelled apical boundaries. We were unable to clearly visualise cortical pMLC at early stages of eye formation (not shown) but immunostaining at the optic cup fusion stage revealed an obvious reduction in the abundance of cortical pMLC in the eyes of *yap1^nl13/nl13^* mutants raised at 32°C (Fig.4A, B; n=12 wt;12 mutant).

**Fig. 4:**
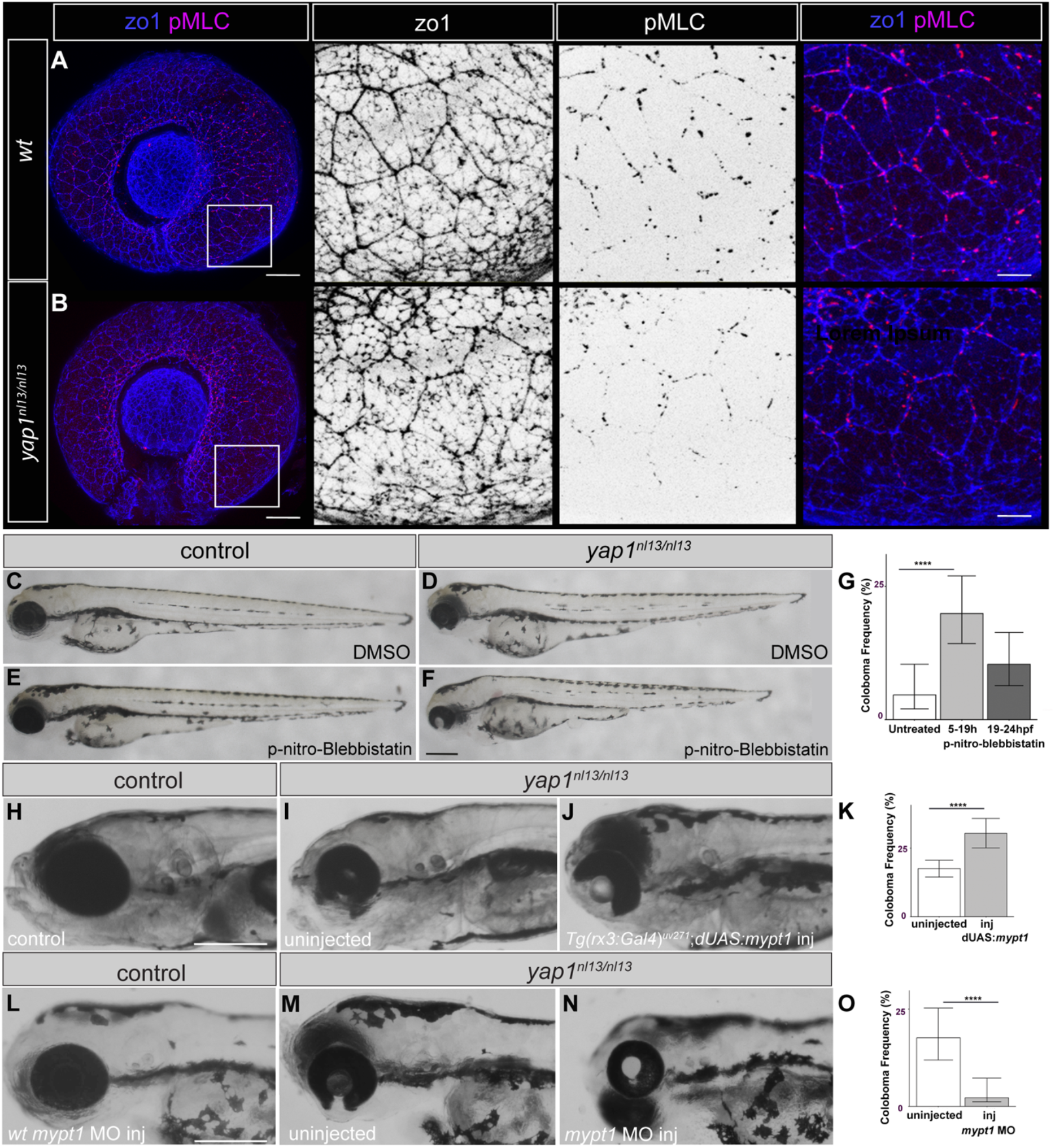
Levels of myosin phosphorylation impact penetrance of coloboma in *yap1^nl13/nl13^* mutants. A,B) 2dpf wildtype and *yap1^nl13/nl13^* mutant eyes showing cell outlines (ZO1 staining) and cortical phosphorylated myosin (pMLC staining) (scale bars, 50µm and 10µm). C-F) Lateral views of 3 dpf wild-type (C,E) and *yap1^nl13/nl13^*mutant (D,F) embryos unexposed (C,D) or exposed (E,F) to 2µM para-nitro- blebbistatin. G) Percentage of embryos showing coloboma in clutches of embryos from *yap1^nl13/+^* x *yap1^nl13/+^* crosses unexposed (control) or exposed to blebbistatin between 5-19 hpf or between 19-24hpf. Note that full penetrance of coloboma in *yap1^nl13/nl13^* mutants would manifest as approximately 25% of embryos in a clutch showing the phenotype. Equality of proportions Qʹ test (*χ*^2^ = 11.4, degrees of freedom = 2, p = 0.003, n = 108, 151 and 145 embryos respectively) and *post hoc* modified Marascuilo procedure with Benjamini-Hochberg correction for multiple testing. H-J) Lateral views of 5 dpf control H) uninjected *yap1^nl13/nl13^* mutant (I) and *Tg(rx3:Gal4)^vu271^*; *UAS:mypt1-RFP*-injected *yap1^nl13/nl13^*mutant (J) embryos. K) Frequency of coloboma in *yap1^nl13/nl13^* mutant embryos expressing *Tg(rx3:Gal4)^vu271^* in uninjected and *UAS:mypt1-RFP*-injected embryos. Fisher’s exact test (p=0.0032, n=598 uninjected and n=283 UAS-injected embryos). -L-N) Lateral views of eyes in wildtype injected with mypt1 MO (L) *yap1^nl13/nl13^*mutant (M), and *yap1^nl13/nl13^* mutant injected with mypt1 MO (N). Scale bar = 200µm. O) Incidence of coloboma in the progeny of *yap1^nl13/+^*incrosses in *mypt1* morphant and control groups. Fisher’s exact test (p=5.5 × 10^-5^, n=125 uninjected embryos, 183 morpholino-injected embryos). Error bars represent 95% confidence interval. **** indicates p ≤ 0.005.

To assess whether reduced actomyosin contractility could contribute to the causation of coloboma in *yap1^nl13/nl13^* mutants, we used blebbistatin to chemically reduce myosin phosphorylation in wildtype and *yap1^nl13/nl13^* embryos. Exposure of wildtype embryos to subthreshold concentration of blebbistatin (2μM) from 5-24hpf had no overt effect on eye development (or on overall development) in wild-type embryos (Fig 4C,E), whereas it led to high penetrance bilateral coloboma in *yap1^nl13/nl13^* mutants raised at 28°C (phenotype occurrence in control clutch, mixed genotype: 5.0±1.3%; treated clutch, mixed genotype: 20.7±1.7% mean ± SEM; Mann-Whitney test p-value = 0.0002; control embryos n=339; treated embryos n=433, image data not shown).

Exposing embryos to blebbistatin for shorter periods from 50% epiboly to the end of the segmentation period (during the 5-24hpf time period), showed that, as for temperature stress, it is the early phase of eye formation that is more sensitive to affecting the later penetrance of coloboma. Thus, *yap1^nl13/nl13^*embryos treated during optic vesicle evagination (50% epiboly to 22s, 5-19hpf) showed high penetrance coloboma while mutants treated from optic vesicle invagination onward (22s-30+ somites; 19-24hpf) showed no significant increase in coloboma penetrance when compared to controls (Fig.4D,F,G).

To assess if it is reduced actomyosin contractility within the eye that affects coloboma penetrance, we overexpressed myosin phosphatase within the forming optic vesicles through injection of dUAS:myr-RFP;*mypt1* plasmid in *Tg(rx3:Gal4-VP16)^vu271^*; *yap1^nl13/nl13^*and sibling embryos. Unlike siblings, *yap1^nl13/nl13^* mutants expressing myosin phosphatase in the optic vesicles showed high penetrance, severe coloboma (Fig.4J) when raised at 28°C (Fig.4J,K). These results confirmed that *yap1^nl13/nl13^* mutants are sensitised to the effects of reducing myosin phosphorylation and likely, as a consequence, actomyosin contractility in the forming eye.

### Reducing myosin phosphatase activity rescues coloboma in *yap1^nl13/nl13^* mutants

Reducing myosin phosphorylation increases penetrance of coloboma in *yap1^nl13/nl13^* mutants; to assess if, in contrast, increasing phosphorylated myosin reduces coloboma penetrance, we used validated morpholinos (MO) to reduce *myosin phosphatase* (*mypt1;* (Gutzman and Sive, 2010)). Injection of *mypt1* MO decreased the penetrance of coloboma in the progeny of *yap1^nl13/+^* in-crosses raised at 30°C (Fig.4L-O) and severity (expressivity) of coloboma when present (Fig.4’M,L). While the coloboma phenotype was less penetrant in *mypt1* MO injected mutants, eyes were smaller when compared with uninjected embryos (Fig.4M,N). Together these data suggest that levels of actomyosin contractility in the forming eye affect the penetrance of coloboma in *yap1^nl13/nl13^* mutants.

### Eye phenotypes are severe when *yap1^nl13/nl13^* and *mab21l2^u517/u517^*mutations are combined

Results above show that the penetrance of coloboma in *yap1^nl13/nl13^*mutants can be increased by environmental stress (higher temperature) and by likely mechanical stress (altered actomyosin activity) in the forming eye. We next assessed if *yap1^nl13/nl13^*phenotypes are also sensitised to the effects of additional mutations in genes/pathways important for eye formation by abrogating a variety of such genes in embryos from in-crosses of *yap1^nl13/+^* fish. Our first focus was on *mab21l2,* a gene than when mutated in humans, fish and mice causes a range of eye phenotypes that overlap with those resulting from compromised *yap1* function (Ceroni et al., 2024; Gath and Gross, 2019; Williamson et al., 2014). In zebrafish, *mab21l2^u517/u517^* mutants show a fully penetrant small eye phenotype and occasional coloboma (Fig.5C, (Wycliffe et al., 2021). Loss of *mab21l2* function in homozygous *yap1^nl13/nl13^*mutants resulted in very severe microphthalmia and fully penetrant bilateral coloboma (Fig.5D; n>200 from multiple crosses) revealing a strong genetic interaction between these two genes.

**Fig. 5:**
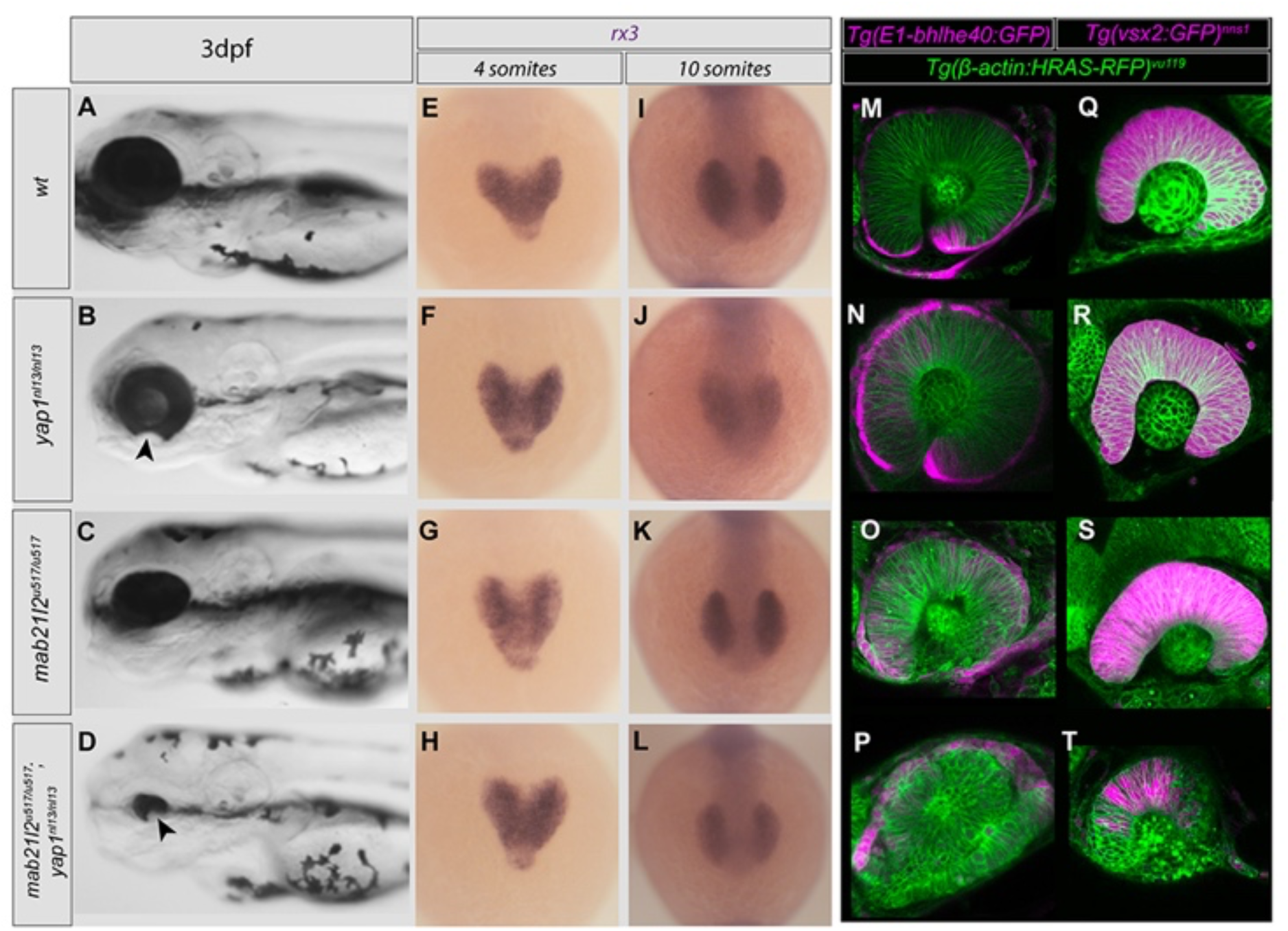
Severe synergistic eye phenotypes upon abrogation of both *yap1^nl13/nl13^* and *mab21l2^u517/u517^.* A-D) Lateral views of 4 dpf wild-type (A), *yap1^nl13/nl13^* (B), *mab21l2^u517/u517^*(C) and *mab21l2^u517/u517^* ;*yap1^nl13/nl13^*embryos (D). Arrowheads indicate coloboma.E-L) Frontal views of *rx3* expression at 4S (E-H) and 10S (I-L) in genotypes indicated in the corresponding row. M-P) Confocal images of RPE (magenta, E1-*bhlhe40*:GFP transgene) in 26hpf embryos with cellular morphologies delineated in green (β-*actin:HRAS-RFP^vu119^* transgene).Q-T) Confocal images of the neural retina (magenta; *vsx2:GFP^nns1^* transgene) in 24hpf embryos with cellular morphologies delineated in green (β-*actin*:*HRAS-RFP^vu119^*transgene).

Although *mab21l2^u517/u517^; yap1^nl13/nl13^* double mutants had severe microphthalmia, the eye field and optic vesicles at early stages were similar size in double and single mutants (Fig.5E-L). However, subsequent to this, the transition from optic vesicle to optic cup was compromised in *mab21l2^u517/u517^; yap1^nl13/nl13^* double mutants with delayed optic cup invagination and RPE flattening compared to single mutants (Fig.5M-T). In double mutants, the RPE did not flatten around the neural retina and RPE cells remained localised dorsally (compare Fig.5P with Fig.5M) as did expression of the neural retina-expressed transgene *vsx2: GFP ^nns1^* (Fig.5T). Fluorescent puncta suggested increased cell death in the lens, ventro-nasal and vento-temporal retina of *mab21l2^u517/u517^*; *yap1^nl13/nl13^*double mutants (compare Fig.5T with 5Q,S), confirmed by TUNEL staining (Fig.S4; compare C with D, and quantification in E).

These data show that in *mab21l2^u517/u517^*; *yap1^nl13/nl13^*double mutants, defects in eye morphogenesis are more pronounced at earlier developmental stages and result in much more severe phenotypes than in single mutants.

### *yap1^nl13/nl13^* eye phenotypes are enhanced by mutations in ECM genes

As the forming eye is embedded within ECM, which in other contexts can modulate Yap function (Dupont et al., 2011; Mamidi et al., 2018), we assessed if compromising ECM genes affected penetrance of *yap1^nl13/nl13^* dependent eye phenotypes. The genes *col9a2, col9a1b* and *lama1* were selected for analysis due to their expression around the developing eye and choroid fissure *(Semina et al., 2006);* Thisse and Thisse https://zfin.org/action/image/view/ZDB-IMAGE-050208-1161#summary; https://zfin.org/action/image/view/ZDB-IMAGE-050208-1161#summary).

While *lama1^nl14/nl14^* mutants showed a partially penetrant (20-40%) and usually unilateral mild coloboma (Fig.6A, B), *lama1^nl14/nl14^;yap1^nl13/nl13^* double mutants had fully penetrant bilateral microphthalmia and coloboma as well as heart oedema and shorter body axes (Fig.6D). While carrying a single *lama1^nl14/+^* mutant allele did not exacerbate the coloboma phenotype in *yap1^nl13/nl13^* mutants (coloboma in n=7/20 *yap1^nl13/nl13^; lama1^nl14^*^/+^ and n=3/12 *yap1^nl13/nl13;^lama1^+/+^* genotyped embryos), the presence of one mutant *yap1^nl13/+^*allele enhanced both penetrance and expressivity of coloboma in *lama1^nl14/nl14^*homozygotes from 23% to 68%.

**Fig 6.**
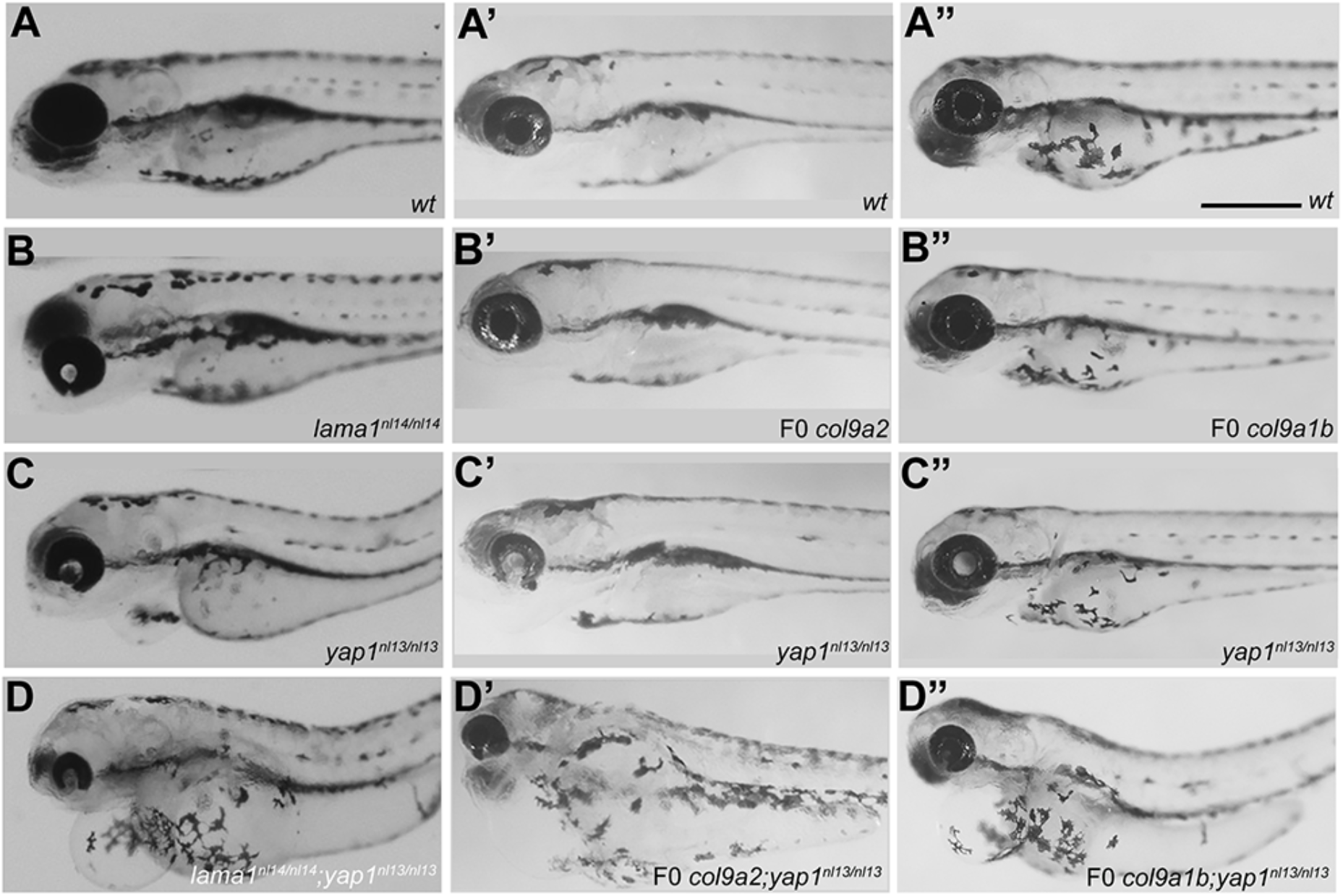
Abrogation of ECM gene function enhances penetrance and severity of *yap1 ^nl13/nl13^*. A-D”) Lateral views, with anterior to the left, of 4dpf embryos with genotypes indicated. A-D) Mild unilateral coloboma shown in (B) was observed in ∼20%-30% *lama1^nl14/nl14^* mutants (n>200; unilateral coloboma in n=3/13 of genotyped *lama1^nl14/nl14^;yap1^+/+^*). Unilateral or bilateral coloboma as shown in C was present in 12% of *yap1^nl13/nl13^* siblings. Severe microphthalmia and bilateral coloboma as shown in D was present in 100% of *lama1^nl14/nl14^/yap1^nl113/nl13^* double mutants (n>200 from several crosses; 9/9 of genotyped double mutants; n=13/19 *lama1^nl14/nl14^*;*yap1^nl113/+^* embryos). A’-D’) No overt phenotype was observed in the eye or elsewhere in F0 *col9a2* crispants (B’; n>100). Unilateral or bilateral coloboma as shown in C’ was present in 7% of the *yap1^nl13/nl13^*siblings (n=12/176). Severe microphthalmia and bilateral coloboma as shown in D’ was present in 27% of F0 *col9a2* injected embryos (n=22/81; genotyping showed that 18/20 of the coloboma group were *yap1^nl13/nl13^* and 2/20 were *yap1^nl13/+^).*A”-D”) No overt phenotype was observed in the eye or elsewhere in the embryo in F0 *col9a1b* crispants (B”; n>120). No coloboma or eye phenotype was observed in *yap1^nl13/nl13^* mutants (as shown in C”; n=0/60). Severe microphthalmia and bilateral coloboma was present in 48% of F0 *col9a1b* injected embryos (as shown in D” n=50/104; genotyping showed that 21/45 of the coloboma group were *yap1^nl13/nl13^* and 24/45 were *yap1^nl13/+^)*.

F0 abrogation of *col9a2* and *col9a1b* function showed no overt phenotypes but abrogation of either gene in *yap1^nl13^* mutants increased penetrance of retinal and other phenotypes (Fig.6A’-D”). Abrogation of *col9a2* increased penetrance and severity of coloboma, microphthalmia and other pleiotropic phenotypes including oedema and shorter body length in *yap1^nl13/nl13^* homozygotes (Fig 6D’). Microphthalmia and other pleiotropic phenotypes was also observed upon abrogation of *col9a1b* in all *yap1^nl13/nl13^* homozygotes (Fig.6D”) and in about 50% of *yap^nl13/nl13^* heterozygotes.

### Genetic interaction screening reveals that eye phenotypes in *yap1^nl13/nl13^* embryos can be modulated by mutations likely to increase Wnt β-catenin levels

The Wnt pathway is implicated in many aspects of eye formation including RPE differentiation and choroid fissure closure (Fuhrmann, 2008; Liu and Nathans, 2008; Westenskow et al., 2009; Zhao et al., 2008). Consequently, we explored if mutations in Wnt pathway genes expressed in the eye affected penetrance and severity of coloboma in *yap1^nl13/nl13^* mutants.

Wnt ligands use FZDs and LRP5/6 coreceptors to initiate Wnt signalling (Tamai et al., 2000) with *fzd5* highly expressed in the developing eye (Cavodeassi et al., 2005; Fuhrmann et al., 2003), *lrp6* expressed widely throughout the CNS and *lrp5* expressed at low levels in the forming RPE and elsewhere in the brain (Fig.S5 A,B). While *lrp5^ulm21/ulm21^* and *lrp6^u348/u348^* mutants showed no overt eye phenotypes (Fig.7E,F; I/J), *fz5^sgu1/sgu1^* mutants showed variable, mild microphthalmia (Fig.7A,B; (Monfries et al., 2025). *lrp6^u348/u348^*/*lrp5^ulm21/ulm21^*double mutants exhibited coloboma, as reported for LRP6 mutant mice, lacked pigmentation and had other phenotypes (Fig.S5G-P, (Zhao et al., 2008), n>200 embryos; no overt phenotype in *lrp6^u348/u348^*/ *lrp5^ulm21/+^* or *lrp6^u348/+^*/ *lrp5^ulm21/ulm21^* embryos) while *fz5^sgu1/sgu1^*;*lrp6^u348/u348^* double mutants were indistinguishable from *fz5^sgu1/sgu1^* single mutants (Fig.S5C,D and data not shown).

**Fig. 7:**
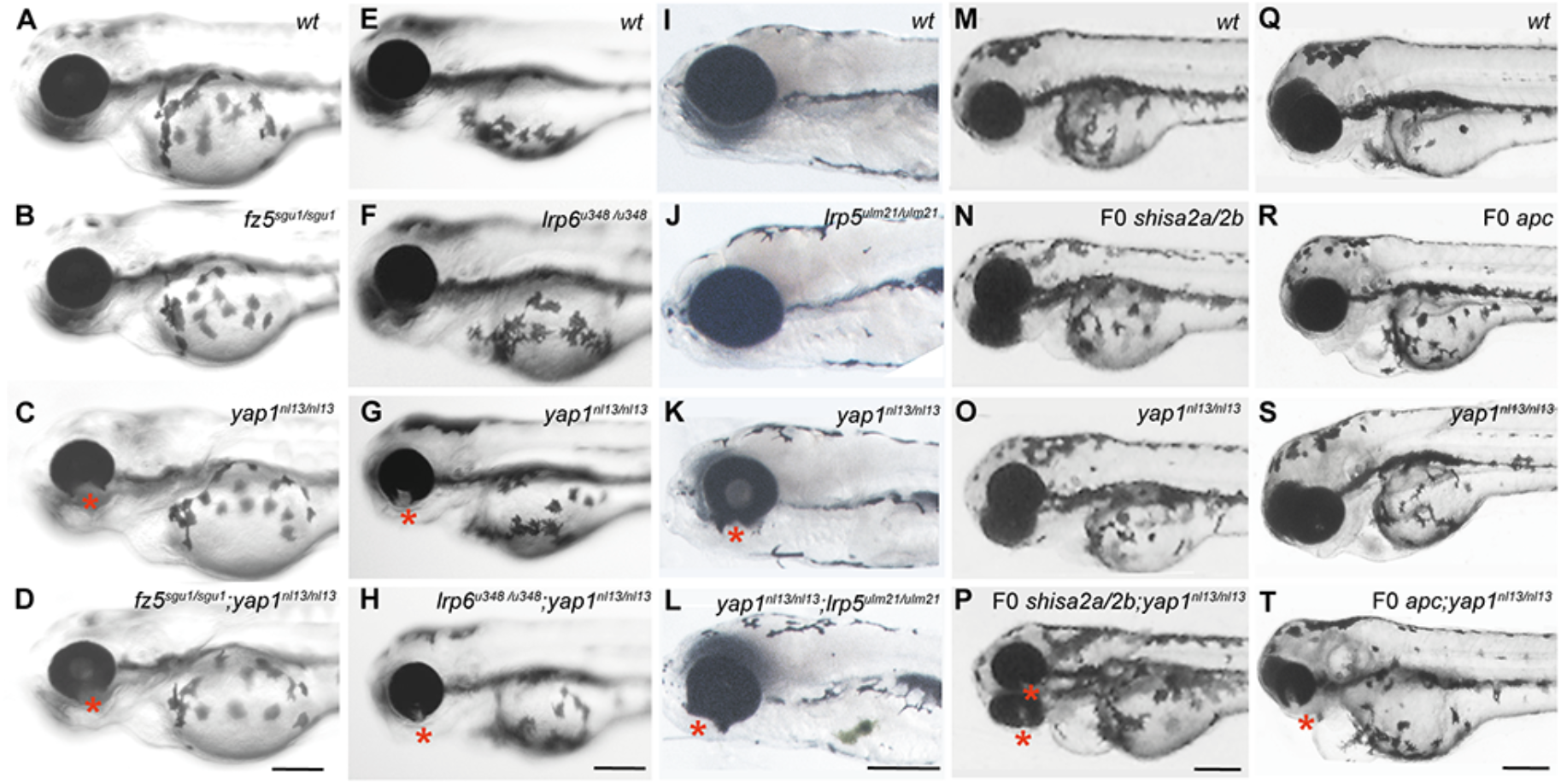
Yap1nl13 shows genetic interaction with shisa2a/2b and apc but not with *fz5, lrp5* or *lrp6.* A-T Lateral views of 3/4 dpf heads with genotypes indicated top right. A-D) Lack of genetic interaction between *fz5^sgu^*^1^ and *yap1^nl13^*. Coloboma was present in 10% of embryos from both single *yap1^nl13/+^* (C; n=80/785) and *yap1^nl13/+^*/*fz5^sgu^*^1/+^ double heterozygous crosses (D, n=81/762). E-H) Lack of genetic interaction between *lrp6^u348^* and *yap1^nl13^*.Coloboma was present in 13.5% of *yap1^nl13/+^* (G, n=26/192) and 11% of *yap1^nl13/+^/lrp6^u348/+^*double heterozygous crosses (H, n=19/250). I-L) Lack of genetic interaction between *lrp5^ulm21^* and *yap1^nl13^*. Coloboma was present in 8% of *yap1^nl13/+^* (K; n=6/75) and 8.5% of *yap1^nl13/+^/lrp5^ulm21/+^*crosses (L; n=13/150). M-P) Genetic interaction between *shisa2a/*2b and *yap1^nl13^*. Coloboma was present in n=14/16 of *yap1^nl13/nl13^*mutants and 3/18 *yap1^nl13/+^* heterozygotes injected with F0 *shisa2a/*2b compare with 3/20 in non-injected *yap1^nl13/nl13^*. Q-T) Genetic interaction between F0 *apc* and *yap1^nl13.^*. Coloboma was observed in n=69/108 of F0 *apc* injected embryos form *yap1^nl13/+^* crosses compare to n=4/67 of non-injected embryos; 32 of the coloboma embryos were genotyped and show no differences in the severity of the phenotype between F0 *apc* injected wildtype and injected *yap1^nl13/nl13^* mutants. Asterisk in C,D,G,H,K,L,P,,T indicate colobomatous eyes. Scale bar = 200µm

Penetrance and expressivity of the *yap1^nl13/nl13^* eye phenotype was not overtly changed in combination with mutations in *fz5^sgu1/sgu1^* (Fig.7 C/D), *lrp6^u348/u348^*(Fig.7G/H) or *lrp5^ulm21/ulm21^*(Fig.7K/L). Indeed, even a small number of analysed *lrp6^u348/u348^*; *fz5^sgu1/sgu1^*; *yap1^nl13/nl13^* triple mutant embryos showed no enhanced eye coloboma or microphthalmia phenotype (Fig.S5 E/F). Consequently, we see no evidence of significant genetic interaction between the *yap1^nl13/nl13^*mutation and mutations in genes encoding Wnt pathway receptors.

As a final approach to altering Wnt receptor function, we abrogated *shisa2a*, a gene expressed in the ventral retina encoding a protein reported to suppress Wnt (and Fgf) signalling by regulating maturation and trafficking of receptor proteins (Liu et al., 2023; Yamamoto et al., 2005); https://zfin.org/ZDB-IMAGE-030925-91#summary;). *shisa2a/2b* crispants did not display any overt phenotype (Fig.7M/N; n>200 from several independent experiments) whereas F0 abrogation of these genes led to almost fully penetrant microphthalmia and coloboma in *yap1^nl13/nl13^* homozygotes (Fig.7O, P). In contrast, there were no enhanced phenotypes upon abrogation of *shisa2a/2b* in either *fz5^sgu1/sgu1^* or *lrp5^ulm21/ulm21^* mutants (Fig.S6A-H).

Given that loss of Shisa2 protein is predicted to increase Wnt signalling, we next assessed the consequences of abrogating *axin1* and *apc*, two widely expressed negative regulators of Wnt signalling with roles in eye formation (Heisenberg et al., 2001; Nadauld et al., 2006). F0 abrogation of *axin1* function in embryos derived from mating *yap1^nl13/+^* heterozygotes led to severe microphthalmia independently of the *yap1* genotype (Fig.S6 I-L).

*apc* crispants showed coloboma and severe microphthalmia when homozygous for *yap1^nl13/nl13^* (Fig.7T). *apc* crispants had no overt morphological eye phenotype in *yap1^+/+^* embryos but showed microphthalmia in about 50% of genotyped *yap1^nl13/+^* heterozygotes (Fig.7R and data not shown). These results were confirmed in embryos carrying both *apc^CA50a/CA50a^* and *yap1^nl13/nl13^* mutant alleles (n=118/526 of embryos derived from *apc^CA50a/+^/yap1^nl13/+^*parents showed microphthalmia and coloboma versus 10/155 of embryos derived from *yap1^nl13/+^* and 0/80 *apc^CA50a/+^* siblings). These interaction phenotypes were suppressed when embryos were raised at 24°C (2/55 with mild microphthalmia in progeny of *apc^CA50a/+^; yap1^nl13/+/^* parents).

## Discussion

In this study, we show that the highly variable penetrance of coloboma in *yap1^nl13/nl13^* mutants is due to genetic and environmental factors influencing the stochasticity inherent in the process of choroid fissure closure. In absence of additional adverse genetic or environmental stressors, the probability that eye morphogenesis proceeds effectively in *yap1^nl13/nl13^* mutants is high; if the process does fail, it is consequently unlikely to do so in both left and right eyes and coloboma is usually unilateral. It is not necessary to postulate a substantive difference of any sort between the developing right and left eyes to explain major differences in final phenotypic outcome. We believe that this interpretation of the causes of phenotypic variability in *yap1^nl13/nl13^* mutants is very likely pertinent to the widely observed variability in many diverse congenital abnormalities of eye formation (Williamson and FitzPatrick, 2014).

### Stochasticity likely underlies phenotypic asymmetries between left and right eyes

While some of the individual variation in congenital eye defects phenotypes in humans is likely explained by differences in allele combinations, why left and right eyes in the same individual can show different phenotypes is less easy to explain. Although it has been speculated that mosaic somatic mutations may contribute to this variability (Chesneau et al., 2023), we consider this to be, in most cases, unlikely. In this study and others , we see comparable variability between left and right eyes in rapidly developing zebrafish embryos in which the likelihood of somatic mutations affecting early development is negligible (Deml et al., 2015; Gestri et al., 2009; Monfries et al., 2025; Shen et al., 2025; Young et al., 2019). Rather, we think it can be misleading to assess the effects of the genetic mutation(s) by correlating genotype solely with the variably penetrant, final eye phenotype. The developmental mechanisms underlying eye formation are robust and can, in some circumstances, cope with deleterious mutations leading to effective recovery from highly penetrant early phenotypes (Young et al., 2019). However, in such circumstances, compensatory mechanisms may fail with variable probability, especially if exposed to additional stressors, thereby leading to differences in penetrance or expressivity of the final eye phenotype. Consistent with this, we observed that eye phenotypes in *yap1^nl13/nl13^* mutants show highly variable penetrance; in absence of additional adverse genetic or environmental factors, coloboma is partially penetrant and usually unilateral whereas in the presence of additional stressors, the penetrance of bilateral coloboma can rise to 100%.

Our interpretation of these observations is that although eye formation can occur normally in *yap1^nl13/nl13^* mutants, the cellular and tissue-level morphogenetic processes that shape the eye are less robust than in wild types. Little is known of the mechanisms that underlie robustness in eye morphogenesis, but in the case of *yap1^nl13/nl13^*mutants, our data suggest that it is the likelihood of mechanical/structural failure in tissue integrity during choroid fissure closure that is increased (as opposed to loss of cells or re-specification of cell types). We find that RPE cells are specified normally and, at least to the level at which our analysis was performed, they flatten, pigment and spread around the eye and into the choroid fissure indistinguishably from wild type. Coloboma appears to result from a stochastically occurring catastrophic failure in the structural integrity of the ventral RPE.

### Yap function in the RPE is required for choroid fissure closure

Coloboma can arise due to disruption in several developmental processes including failure to appose the nasal and temporal lips of the choroid fissure, failure to degrade the extracellular matrix within the fissures leading to failure in epithelial fusion, or re-specification of retinal cell types (Gestri et al., 2018; James et al., 2016; Yoon et al., 2020). Consequently, disrupted function in genes functioning both in ventral neural retina and in the periocular mesenchyme that invests the eye can lead to coloboma and this study additionally demonstrates the importance RPE cells in fissure closure (Lupo et al., 2011). Although *yap1* is expressed and presumably functions in many cell types in and around the eye (Kim et al., 2016; Miesfeld and Link, 2014; Williamson et al., 2014), we show that restoration of wildtype Yap1 function solely within the RPE suppresses the coloboma phenotype. Disruption of several other RPE-expressed genes, including *fat1* and *nlz2/zfp503*, leads to coloboma (Boobalan et al., 2022; Lahrouchi et al., 2019) but the broad expression of such genes has precluded conclusions on tissue specificity of their requirement for fissure fusion. Loss of ß- catenin function in the RPE leads to trans-differentiation of RPE to neural retina and this is associated with a variety of retinal morphogenesis defects including coloboma but the severity of these defects does not allow conclusions about the normal role of the RPE in fissure closure (Westenskow et al., 2009). More directly pertinent is a recent study suggesting that loss of pigmentation and trans-differentiation of cells lining the choroid fissure are causative of coloboma in zebrafish carrying mutations in *yap1^bns19/bns91^* plus *taz^+/-^* (Neelathi et al., 2026). Conversely, our study shows that coloboma can occur in absence of the more severe ventral retinal deficits evident in *yap1^bns19/bns91^; taz^+/-^* mutants when, morphologically, the *yap1^nl13/nl13^* mutant RPE is indistinguishable from wildtype prior to rupture. However, it is intriguing that the cells lining the fissure derive from the RPE and it is, of course, possible that the structural deficits in the RPE that we propose to underlie coloboma could, at least in part, be impacted by altered expression of genes that confer RPE identity (Buono et al., 2021).

### Impaired myosin-mediated contractility in *yap1*^nl13/nl13^ RPE cells is likely to underlie the failure in choroid fissure closure

The implication of RPE cell function in choroid fissure closure is consistent with the suggested role for the RPE in constraining the shape of the developing optic vesicle. It has been proposed that accumulation of phosphorylated myosin in the RPE generates higher stiffness compared to the neural retina leading to invagination and optic cup formation (Carpenter et al., 2015; Eiraku et al., 2011; Moreno-Mármol et al., 2021).

Yap mediated regulation of mechanical force and tissue tension is a feature of many developmental processes with Yap signalling acting as a mechano-transducer of such tissue stress (Panciera et al., 2017; Porazinski et al., 2015). Our data is consistent with changes in the mechanical properties of the RPE in *yap1^nl13/nl13^*eyes being causative of the variably penetrant coloboma phenotypes. RPE cell specification and expansion are not affected in *yap1^nl13/nl13^* mutants; instead, we observed coloboma to arise from rupture of the RPE in the ventral retina as the lips of the choroid fissure appose suggesting this location at this time is particularly prone to mechanical stress. At present, we do not have any direct measures of localised tissue tension but our experiments manipulating levels of phosphorylated myosin support the idea that actomyosin contractility within the RPE is impaired, potentially weakening the ability of the highly stretched RPE cells to contain the neural retina. Notably, we found that pharmacological or genetic approaches that enhance the levels of phosphorylated myosin suppress the occurrence of coloboma, presumably by increasing mechanical robustness.

### Disruptions to *mab21l2* and genes encoding ECM proteins investing the eye enhance phenotypic penetrance and severity

In addition to actomyosin contractility, extracellular matrix (ECM) deposition is a critical factor in regulating tissue stiffness and it is thought that the ECM provides a dynamic and flexible scaffold for many morphogenetic processes (Díaz-de-la-Loza and Stramer, 2024). Indeed, mutations affecting both ECM genes and molecules such as Integrins and Paxillins anchoring the neuroepithelium to the ECM frequently exhibit eye morphogenesis defects (Bryan et al., 2020, 2016; Eintracht et al., 2024; Martinez-Morales et al., 2009; Semina et al., 2006) and ECM components are required for eye organoids to acquire a cup shape (Bazin-Lopez et al., 2015; Eiraku et al., 2011). Consistent with this, we observed strong genetic interactions between *yap1^nl13/nl13^* and mutations in *laminin* and *collagen* ECM encoding genes affecting both penetrance and severity of eye phenotypes. An interplay between Yap and ECM has been described in various other developmental contexts with Yap activity believed to be directly regulated by ECM stiffness (Calvo et al., 2013; Chaudhuri et al., 2014; Dupont, 2016; Dupont et al., 2011). In *yap1^nl13/nl13^*mutants, we suggest that the concurrent loss of structural ECM components like *lama1* compromises an external buffer of mechanical stress. Without this physical scaffolding to constrain the forming optic cup, the structural integrity of the ventral optic cup is more prone to failure, leading to tissue collapse and coloboma. Our RNAseq data also suggest that disrupted yap1 function reciprocally regulates ECM genes with several such genes including *ccn2a* and *ccn1l1* (Fernando et al., 2010)*, serpinh1b* (Kudoh et al., 2001), *fn1b* (Baxendale et al., 2009) and *clusterin* (Jiao et al., 2011) showing altered levels of expression in *yap1^nl13/nl13^* mutants.

We also observed a strong genetic interaction between *yap1* and *mab21l2*, a gene encoding a protein with RNA binding activity of unknown function that, like *yap1*, can be causative of coloboma in humans when disrupted (Ceroni et al., 2024; Deml et al., 2015; Rainger et al., 2014). In zebrafish fully lacking *mab21l2* function, optic vesicle morphogenesis is more severely disrupted than in *yap1* mutants (Deml et al., 2015; Gath and Gross, 2019; Wycliffe et al., 2021). *yap1^nl13/nl13^*embryos lacking *mab21l2* show severe microphthalmia and coloboma; this reveals that although *yap1^nl13/nl13^* mutant eyes can undergo normal morphogenesis up to the stage of fissure fusion, compromised Yap1 function during early phases of eye formation can have phenotypic consequences upon morphogenesis when *mab21l2* function is absent.

Yap can synergise with, or antagonize, Wnt signaling and modulate both canonical Wnt/ßcatenin and non-canonical signalling (Astone et al., 2024; Azzolin et al., 2014; Park et al., 2015; Totaro et al., 2018). Furthermore both *yap1* expression and Wnt/β- catenin signalling are detected in the RPE across multiple vertebrate species (Cho and Cepko, 2006; Dorsky et al., 2001; Miesfeld and Link, 2014; Van Raay et al., 2005; Veien et al., 2008; Wang et al., 2018; Williamson et al., 2014) and disruption to Wnt and Yap/Taz pathways result in similar phenotypes (Fujimura et al., 2009; Holt et al., 2022; Kim et al., 2016; Lieven and Rüther, 2011; Liu and Nathans, 2008; Miesfeld and Link, 2014; Monfries et al., 2025; Zhao et al., 2008). Given these results, we were surprised that we did not see genetic interactions when abrogating various Wnt pathway genes in *yap1*^nl13/nl13^ mutants nor did we observe significant changes in Wnt pathway gene expression in *yap1*^nl13/nl13^ mutants. We did observe genetic interactions when either *shisa2a/2b* or *apc* were reduced/abrogated in *yap1^nl13/nl13^* mutants; although neither of these genes function only within the Wnt pathway (Nadauld et al., 2006; Nagano et al., 2006), Wnt activity is likely increased upon their abrogation raising the possibility of a genetic interaction with *yap1*^nl13/nl13^ only when the Wnt pathway is overactivated.

### Early developmental stressors can have delayed phenotypic consequences

We found that adverse phenotypic modulators (heat or manipulations affecting myosin phosphorylation) impacted penetrance of coloboma in *yap1*^nl13/nl13^ eyes long before the overt failure in choroid fissure closure; this suggests that the robustness of eye morphogenesis is compromised and sensitised to additional stressors from the stage of early optic vesicle formation even though phenotypic consequences only occur much later; that is, there is a “hidden” phenotypic trait in *yap1*^nl13/nl13^ eyes that variably leads to a later overt phenotype. Analogous to this, we have previously found that *tcf7l1a* mutants have reduced eye-field size that can lead to complete loss of eyes or have no consequence on eye formation dependent on genetic background (Young et al., 2019). In the case of *yap1*^nl13/nl13^ eyes, given that manipulating acto-myosin contractility impacts the likelihood of occurrence of the overt coloboma phenotype, it is likely that the covert, hidden, trait is a cytoskeletal change that impacts the structural robustness of RPE cells as they undergo highly dynamic remodelling and movements during optic vesicle formation.

The major differences in phenotypic outcome dependent on exogenous stressors in *yap1*^nl13/nl13^ eyes reveals the plasticity inherent in developmental processes (Metcalfe, 2024). There are numerous examples of developmental trajectories that vary dependent on environmental conditions, often revealing both the robustness built into developmental pathways and the fragility and/or costs of compensatory processes. For instance, stickleback fish that undergo rapid “catch-up” growth after a period of adverse environmental conditions show poor performance in many subsequent tasks reflecting the hidden cost of the initial robust, compensatory response to the developmental challenge (Lee et al., 2013). Our studies show that the robustness inherent in the developmental processes underlying eye development usually enables normal eyes to form in *yap1*^nl13/nl13^ mutants but that such eyes are compromised in their ability to compensate the effects of any additional adverse genetic or environmental stressors.

## Summary

Our study reveals that the RPE plays a critical role in the closure of the choroid fissure most likely through its structural integrity constraining the morphogenesis of the closing optic cup. We show that although eye formation can occur normally in *yap1*^nl13/nl13^ eyes, such eyes are sensitised to the effects of any other environmental, exogenous, or genetic stressors. By studying differences between left and right eyes, our study has revealed that widely different phenotypes can arise despite developing tissues of the same genotype being exposed to the same environment – simply through the stochasticity inherent in developmental outcomes. We suspect such stochasticity to be a likely contributor to the widespread occurrence of phenotypic variability in congenital abnormalities of eye formation.

## Acknowledgments

We are very grateful to members of our group for discussion and suggestions and our colleague Alessandro Mongera for critically reading the manuscript. We thank the Image Facility team for their support with image acquisition and Heather Callaway and the Zebrafish Facility team at UCL, for their fish care.

## Author contributions

N.F., M.S., G.C., L.T., A.M., E.H., S.C., L.E.V., R.Y., M.T., G.G. performed the experiments. D.W., G.W., F.C., and A.N. generated zebrafish lines. N.F. and G.P. performed statistical analyses. R.J.P performed RNA-seq analysis. K.T. and G.G. prepared the figures. S.W.W. and GG wrote the manuscript. All authors read, revised, and edited the manuscript.

## Supplementary Figures

**Fig. S1:**
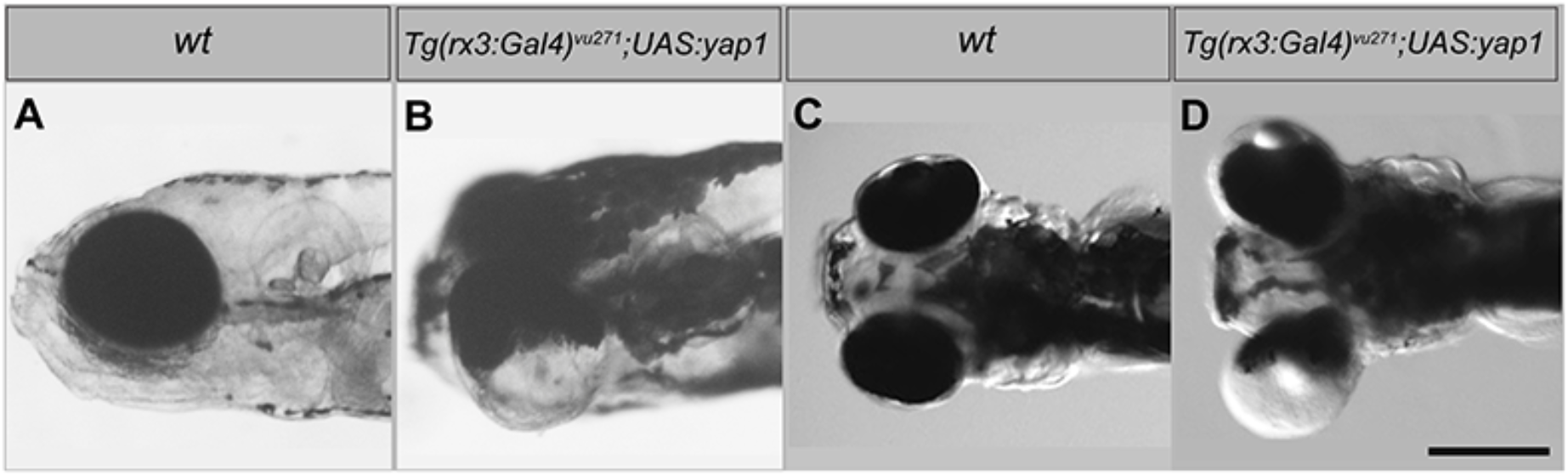
Expressing exogenous Yap1 in the developing neural retina causes severe retinal defects. Lateral (A,B) and dorsal (C,D) views of 4dpf wildtype embryos (A and C) and wild-type embryos overexpressing *yap1* using an *Tg*(*rx3:Gal4)^vu271Tg^* driver (B and D).

**Fig. S2:**
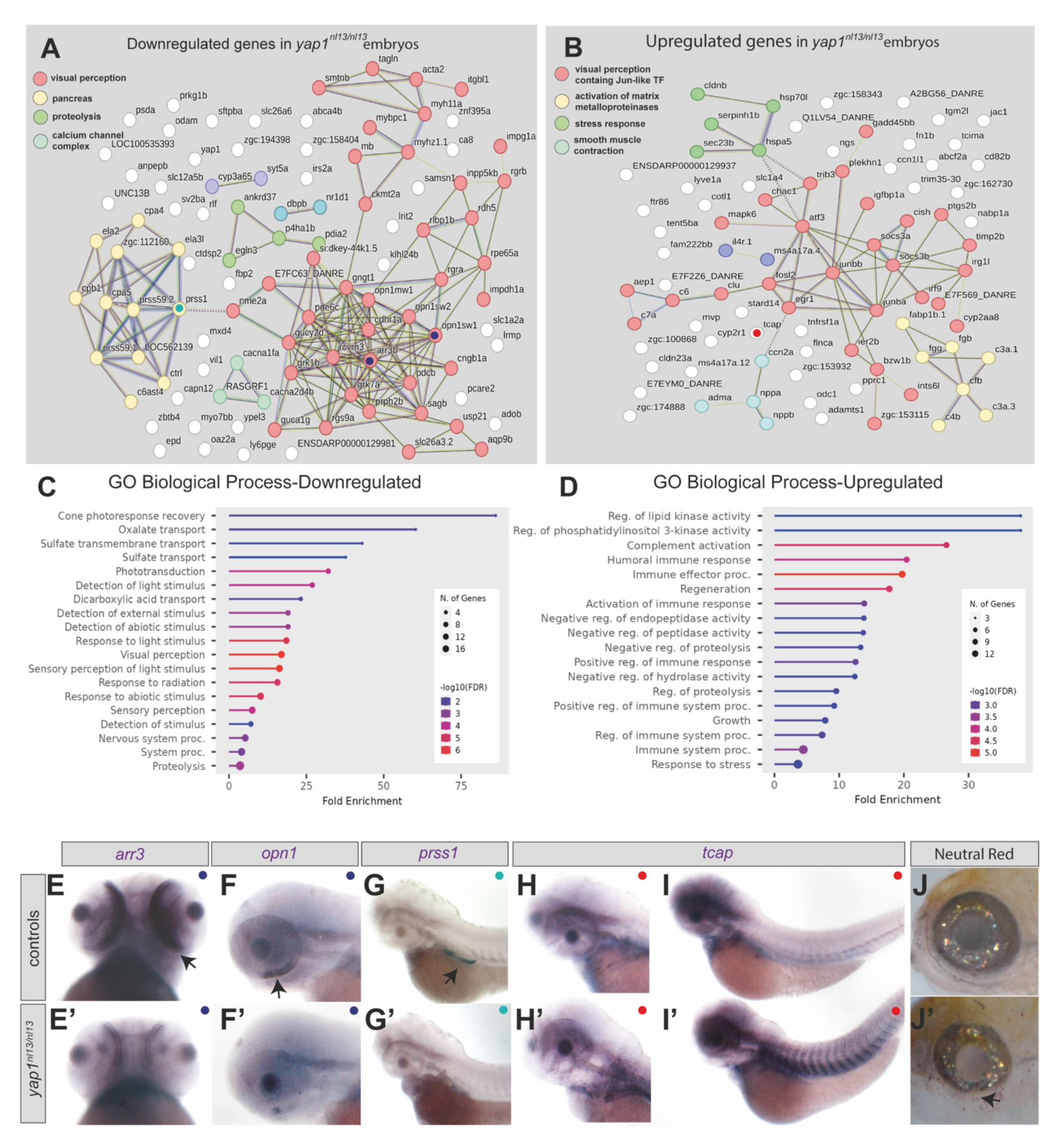
Outer retinal and stress response genes are mis-regulated in *yap1^nl13/nl13^*mutants. A,B) STRING analysis of proteins encoded by genes downregulated (A) or upregulated (B) in 3dpf *yap1^nl13/nl13^* mutants from RNAseq analysis of wild-type and *yap1^nl13/nl13^*embryos with colobomatous eyes raised at standard temperature (28°C). 38% of downregulated and 30% of upregulated genes were expressed in the eye (“visual perception” pink dots; 40/104 and 27/89 respectively) and 11.5% of downregulated and 8% of upregulated genes were expressed in the pancreas (“pancreas” yellow dots in A and table1; 12/104 and 7/89 respectively) based on a q value of <0.1 (See Table1). Many of the downregulated genes were associated with the visual cycle including *arr3* and *RPE65a*, which are expressed in the RPE, and several opsin genes expressed in photoreceptors. In contrast to the downregulation of outer retinal genes, immune response and Jun transcription network genes (yellow dots and a subset of pink dots respectively in (B) implicated in cellular responses to stress and stress response genes (green dots) were upregulated. C,D) GO term analysis of genes with altered expression in *yap1^nl13/nl13^*mutants. E-I’) Eyes of 3dpf embryos (C-C’, ventral views; D-G’, lateral views) showing examples of expression of genes (bottom right in each panel) downregulated in the outer retina (C-D’) or pancreas (E, E’) or upregulated in the ventral somites and elsewhere (F-G’). Coloured dots in E-I’ match locations of genes in clusters in panels A and B. J-J’) Neutral red labelled cells, likely macrophages, around the eye in a *yap1^nl13/nl13^*mutant.

**Fig. S3:**
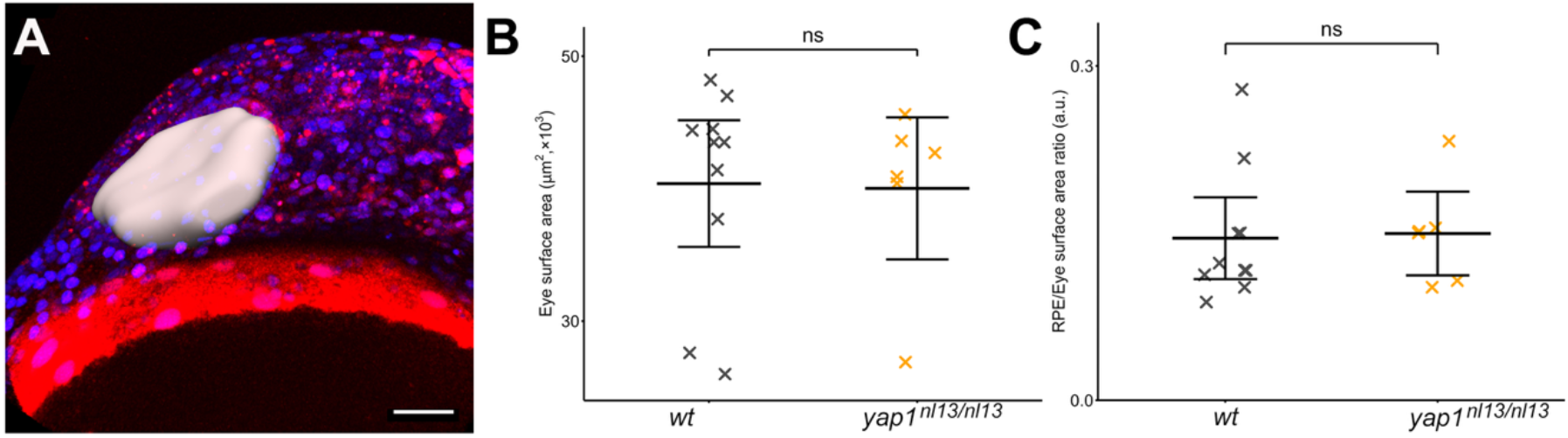
No differences in optic vesicle size and RPE coverage of the optic vesicle between controls and *yap1^nl13/nl13^* mutants. A) Lateral view of the head of a 6s stage embryo illustrating the mask used to estimate the size of the prospective optic vesicle. B) Graph plotting surface area of optic vesicles. At the stage of RPE specification (8/10s), the eyes of wild types and *yap1^nl13/nl13^* mutants have no significant difference in overall surface (eye surface: wild type = 403.8 ± 24.4 μm, N=10; *yap1^nl13/nl13^*= 400.2 ± 27.34, N=6; Mean ± SEM; unpaired *t*-test, p=0.93). C) Graph plotting the ratio between RPE surface and overall eye surface, representing the overall RPE coverage of the eye. There is no significant difference between wild type and *yap1^nl13/nl13^* embryos (RPE Surface /Eye surface ratio: Wild Type= 0.145 ± 0.02 N=10 *yap1^nl13/nl13^* 0.149 ± 0.02 N=6 0.9 Mean±SEM; unpaired t-test for unpaired data p=0.88). Scale bar 50 μM.

**Fig. S4:**
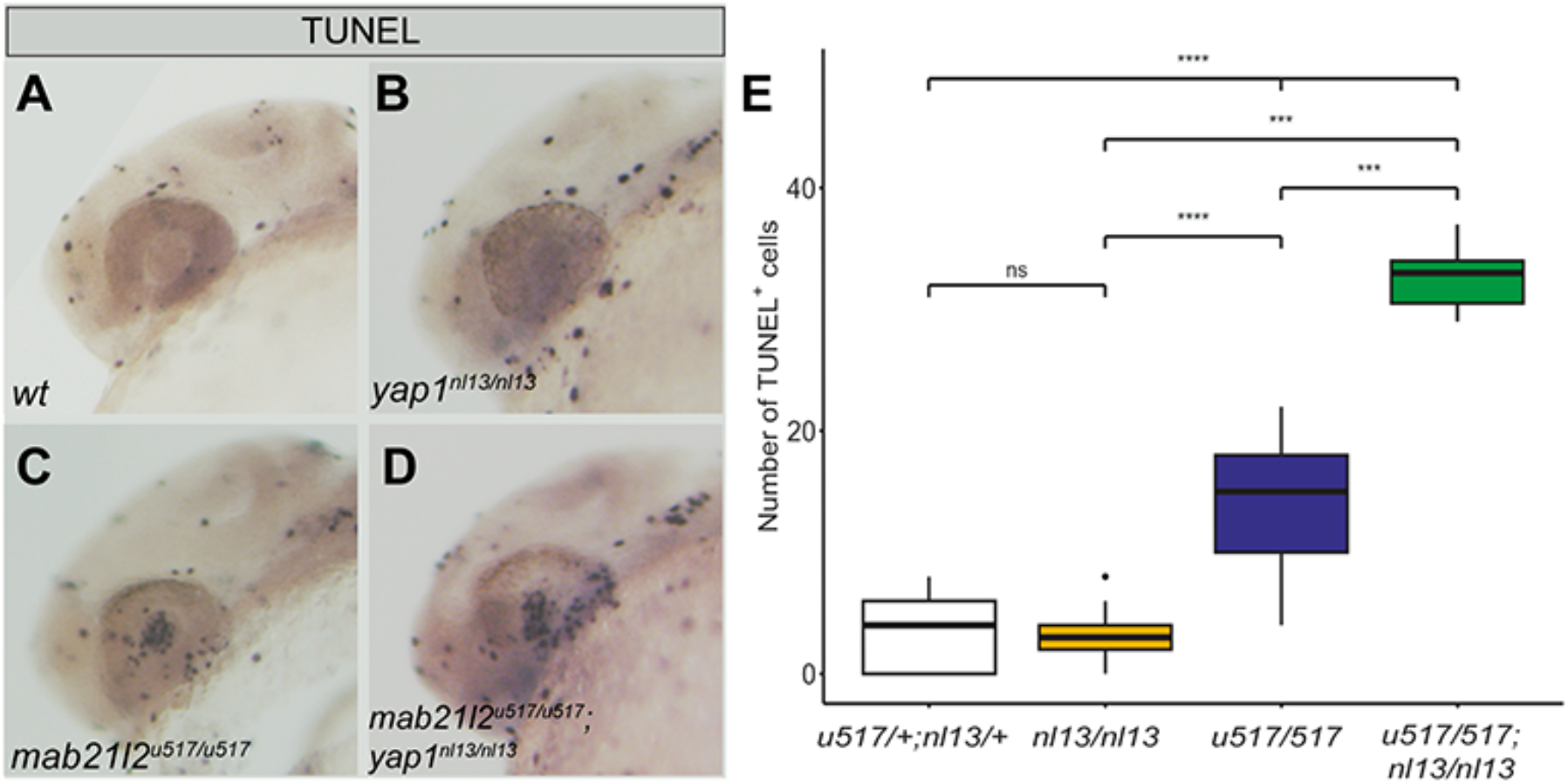
Increased cell death in/around *mab21l2^u517/u517^;yap1^nl13/nl13^* eyes. A-D) Lateral views of 26hpf eyes showing apoptotic (TUNEL+) cells in each genotype shown. E) Numbers of apoptotic cells in the eyes of a 26hpf stage embryos of genotypes shown on the x-axis. Mean ± 95% confidence interval: double heterozygous siblings = 3.5 ± 0.6, *yap1^nl13/nl13^* embryos = 3.0 ± 1.0, *mab21l2^u517/u517^* = 14.6 ± 2.0, *mab21l2^u517/u517^*, *yap1^nl13/nl13^* double mutants = 32.7 ± 2.4. n = 73, 18, 25 and 6 embryos respectively. Kruskal-Wallis test, χ^2^ = 64.8, degrees of freedom = 3, p = 5.49 × 10^-14^, followed by post hoc pairwise comparisons using Wilcoxon rank sum test with continuity correction and Benjamini-Hochberg adjustment for multiple testing. Selected comparisons: *yap1^nl13/nl13^*embryos v. *mab21l2^u517/u517^*, *yap1^nl13/nl13^* double mutants, p = 3.7 × 10^-4^; *mab21l2^u517/u517^* embryos v. *mab21l2^u517/u517^*, *yap1^nl13/nl13^* double mutants, p = 2.6 × 10^-4^.

**Fig. S5.**
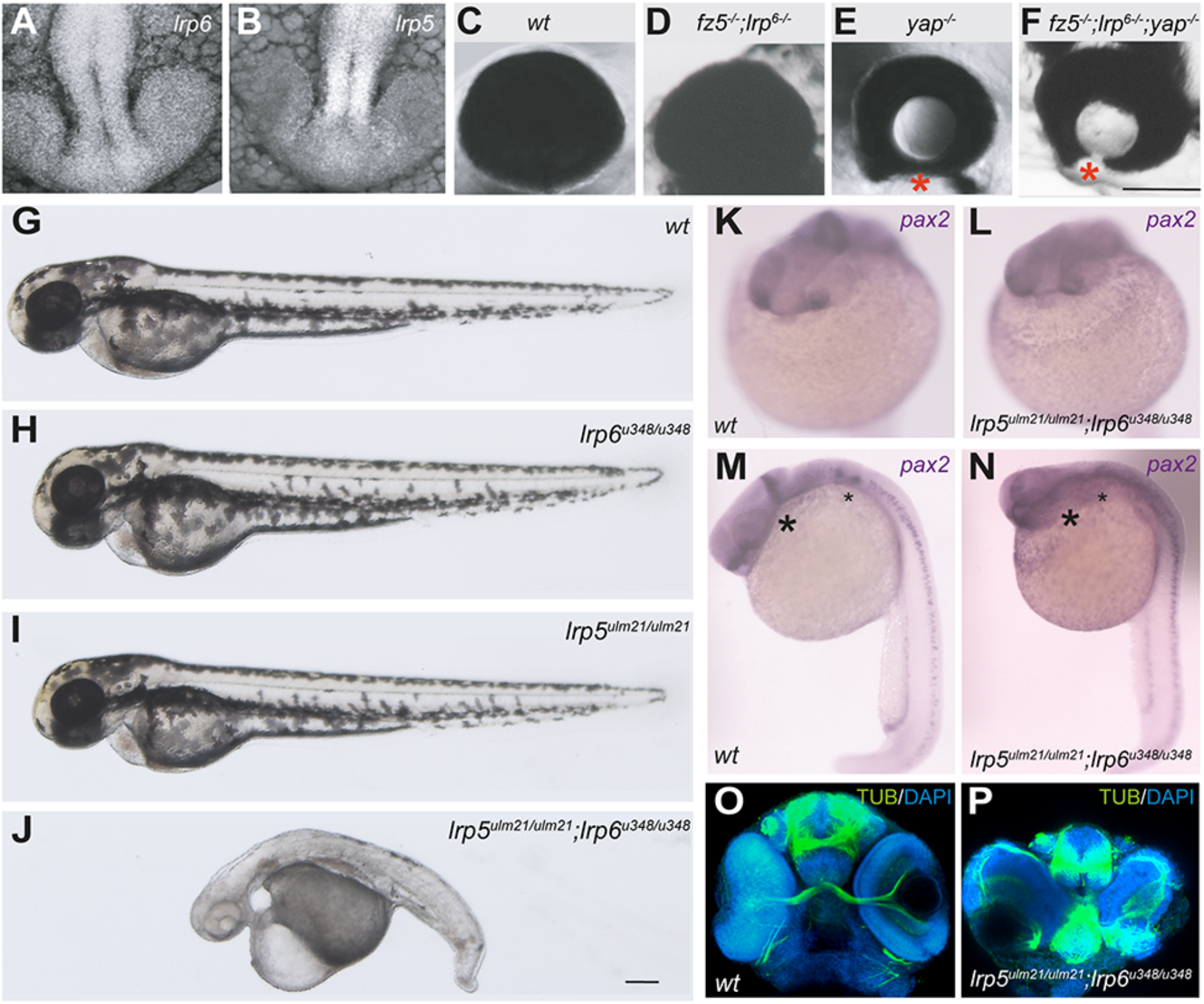
Eye phenotypes in embryos carrying mutations in Wnt pathway genes. A,B) Dorsal view with anterior to the bottom of brains and eyes of 22hpf embryos showing expression of *lrp6* (A) and *lrp5* (B) using HCR. C-F) Lack of genetic interaction between *fz5^sgu1^*;*lrp6^u348^ and yap1^nl13^*. 3dpf eyes of wildtype (C) *fz5^sgu1/sgu1^*;*lrp6^u348/u348^* (D) *yap1^nl13/nl13^* (E) and *fz5^sgu1/sgu1^*;*lrp6^u348/u348^;yap1^nl13/nl13^* (F).Coloboma was present in 13.5% of embryos from *fz5^sgu1/+^*;*lrp6^u348/+^;yap1^nl13/+^*crosses (n=148/1109) versus 14% in *yap1^nl13/nl13^* siblings (n=29/215); from 192 genotyped embryos we found three triple mutants, 1 with bilateral coloboma. Not all allele identifiers are shown. G-J) Lateral views of 2 dpf embryos with genotypes indicated in the top right of each panel. 100% of *lrp5^ulm21/ulm21^;lrp6^u348/u348^*double mutant show lack of pigmentation, shorten body axes and heart oedema as shown in (J). K-N) 22hpf embryos showing expression of *pax2* in the optic stalk (K), retained in the *lrp5^ulm21/ulm21^;lrp6^u348/u348^*double mutant (L) but reduced in the midbrain hindbrain boundary (asterisk in M,N) and rhombomeres compared to wildtypes. Fronto-lateral views (K,L) and laterals view (M,N) with anterior to the left. O/P) Acetylated α-tubulin labelling of axons in wildtype (O) and *lrp5^ulm21/ulm21^;lrp6^u348/u348^*double mutant (P) 48hpf embryos. Frontal views showing severe axonal defects in the eyes and brain of the mutant. Scale bar = 200µm

**Fig. S6.**
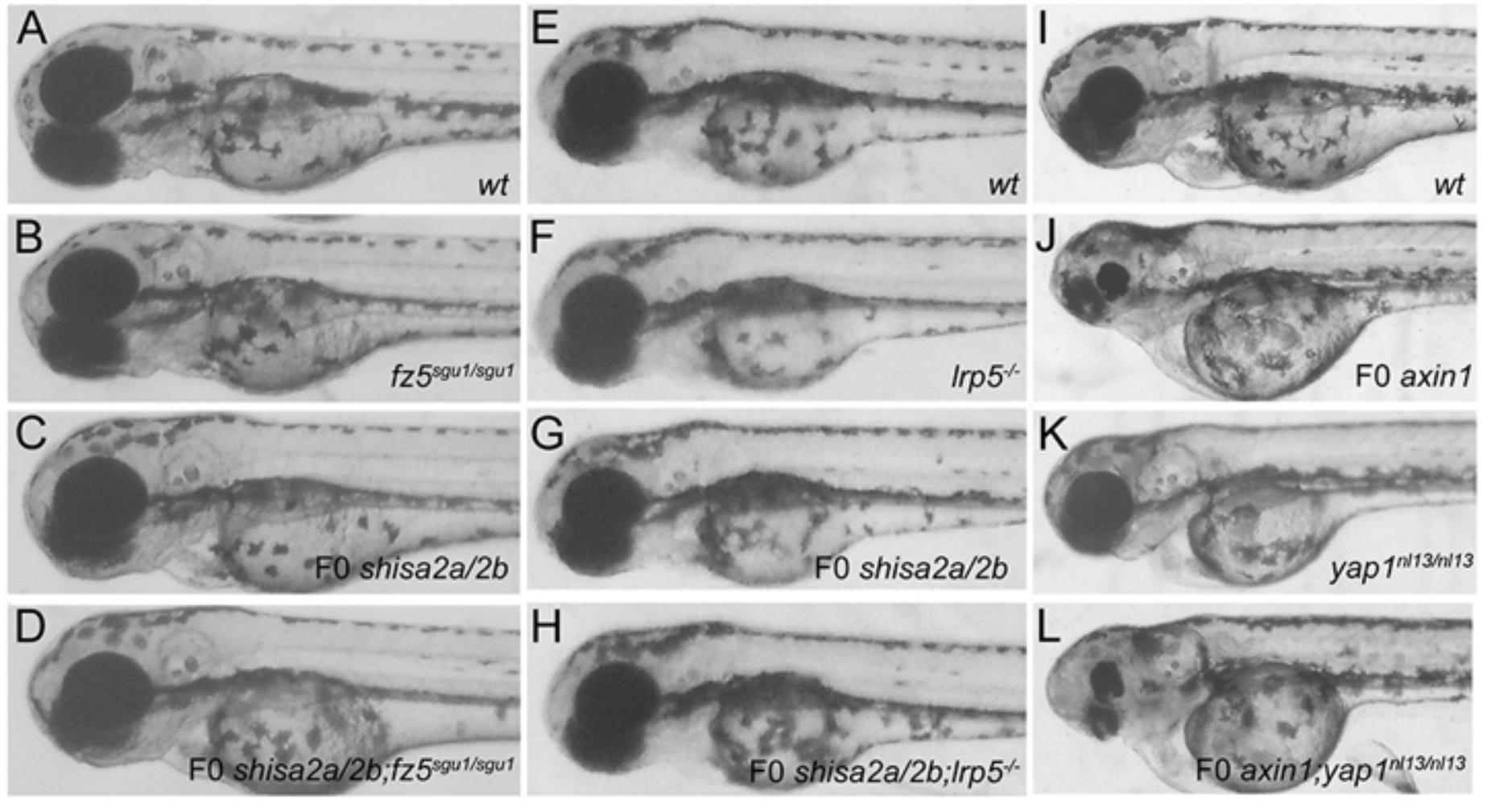
Eye phenotypes in embryos carrying mutations in Wnt pathway genes. A-L) Lateral views of 3-4 dpf heads with genotypes indicated. A-H) No enhanced phenotypes were observed upon abrogation of F0 *shisa2a/2b* in either *fz5^sgu1/sgu1^* (D) or *lrp5^ulm21/ulm21^* mutants (H; 0/140 embryos from *fzd5^sgu/+^* incrosses; 0/88 embryos from *lrp5^ulm21/+^* incrosses). I-L) F0 abrogation of F0 *axin1* function gave rise to severe microphthalmia in wildtype (J) and *yap1^nl13^* mutants (L).

## Supplementary Tables

**Table S1:**
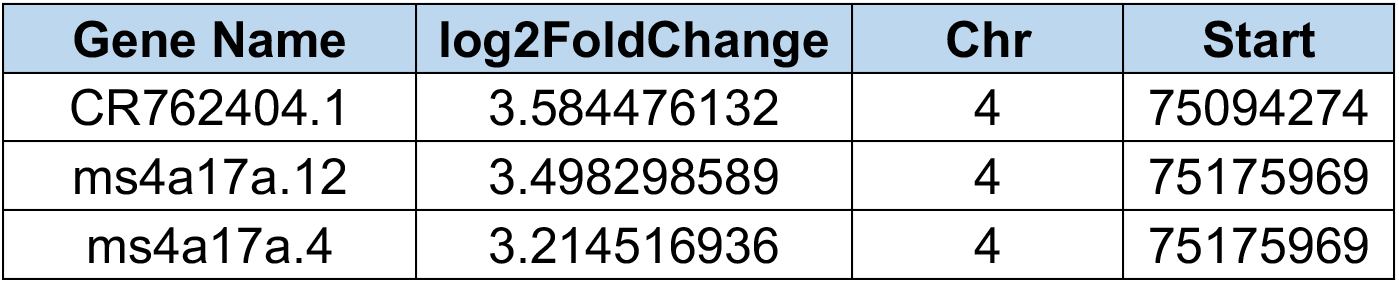
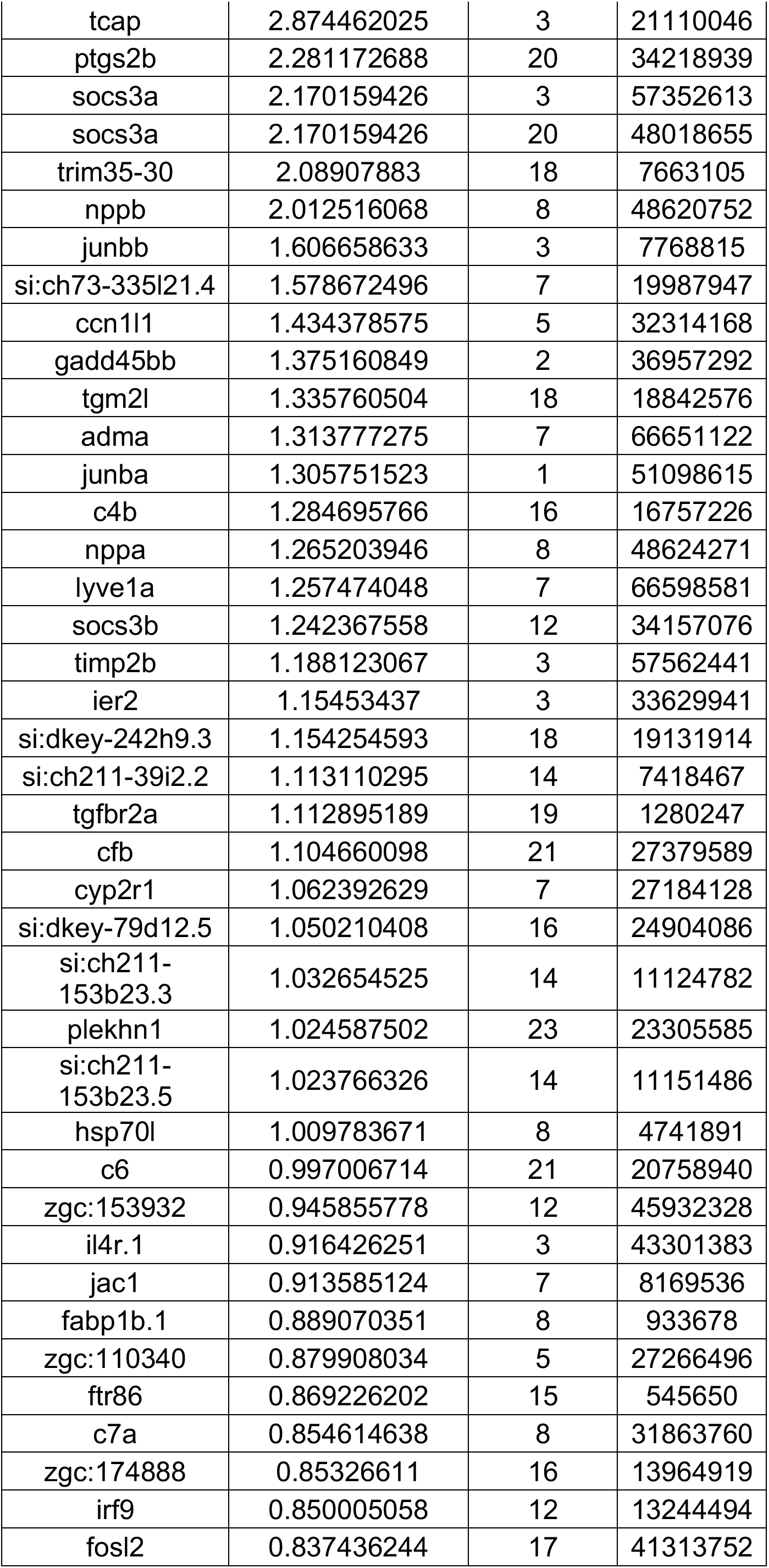
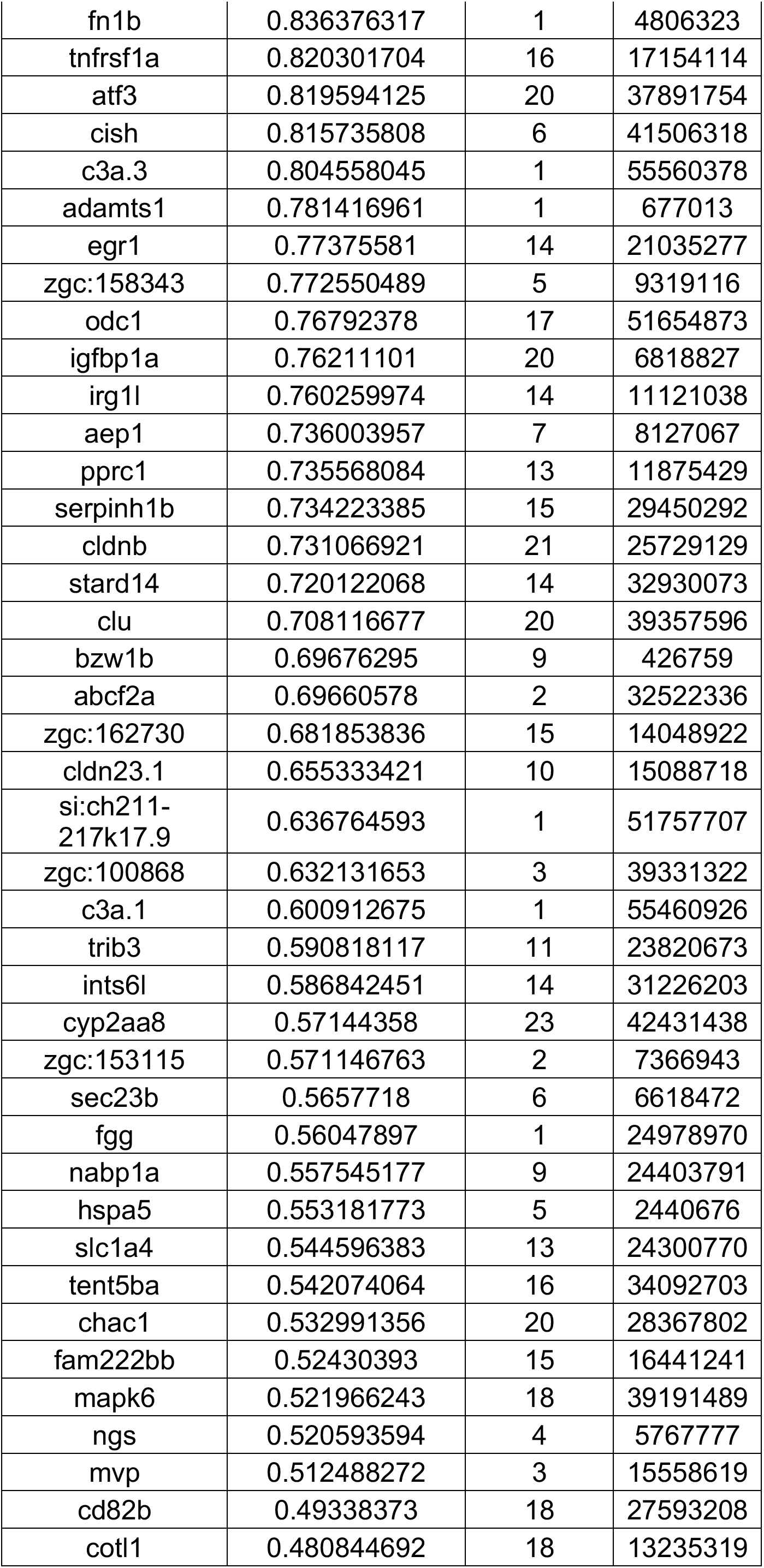
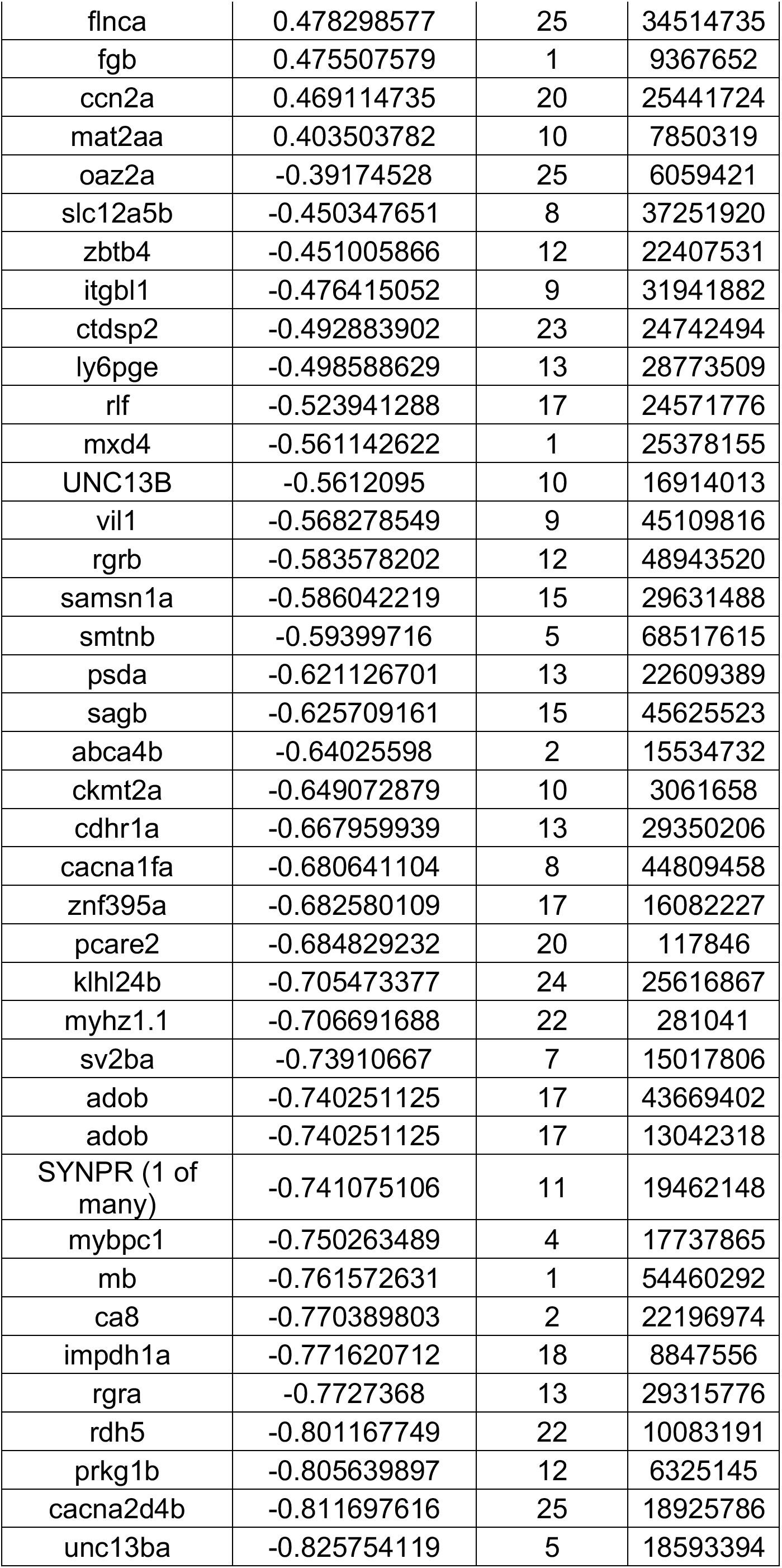
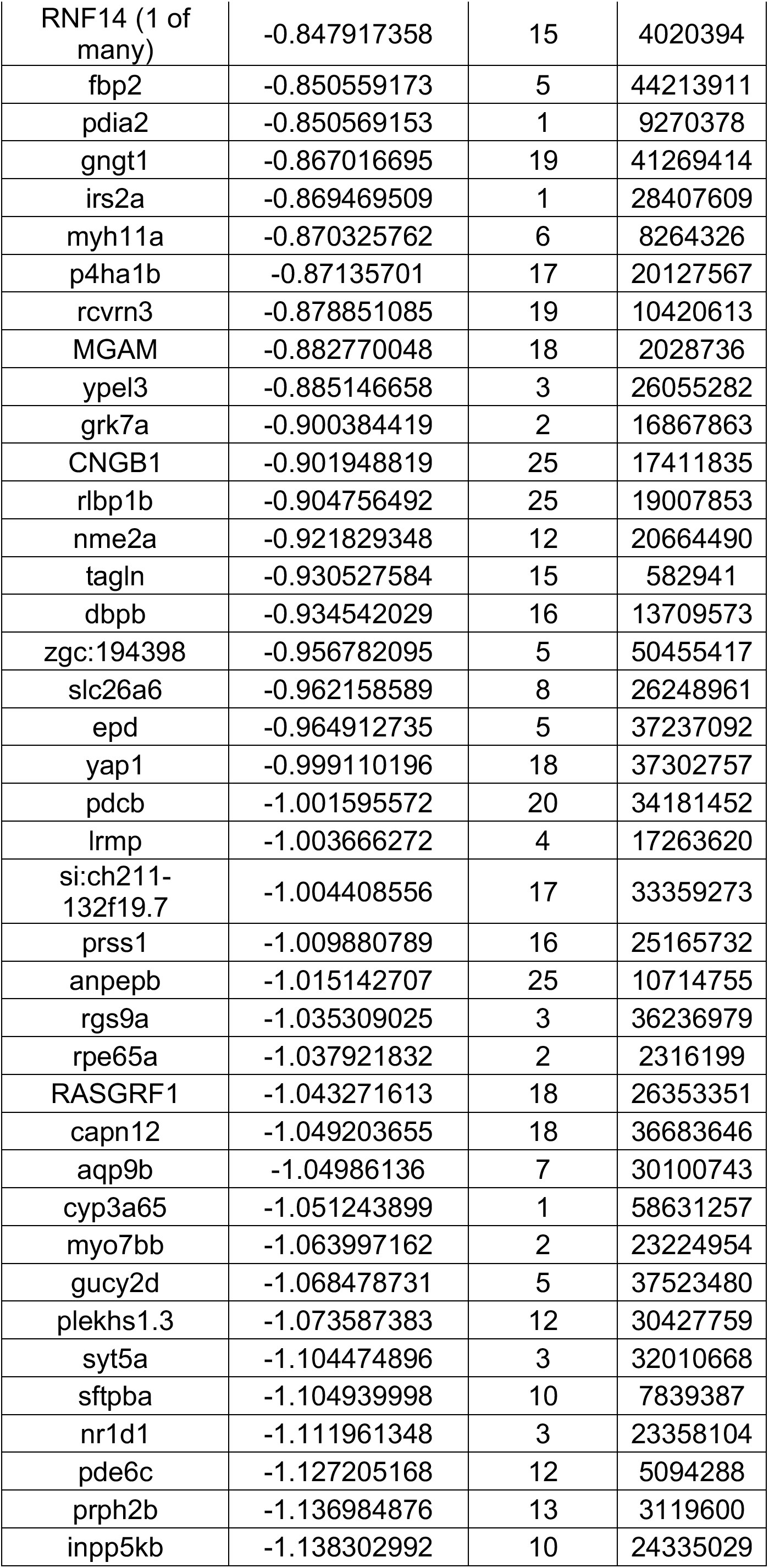
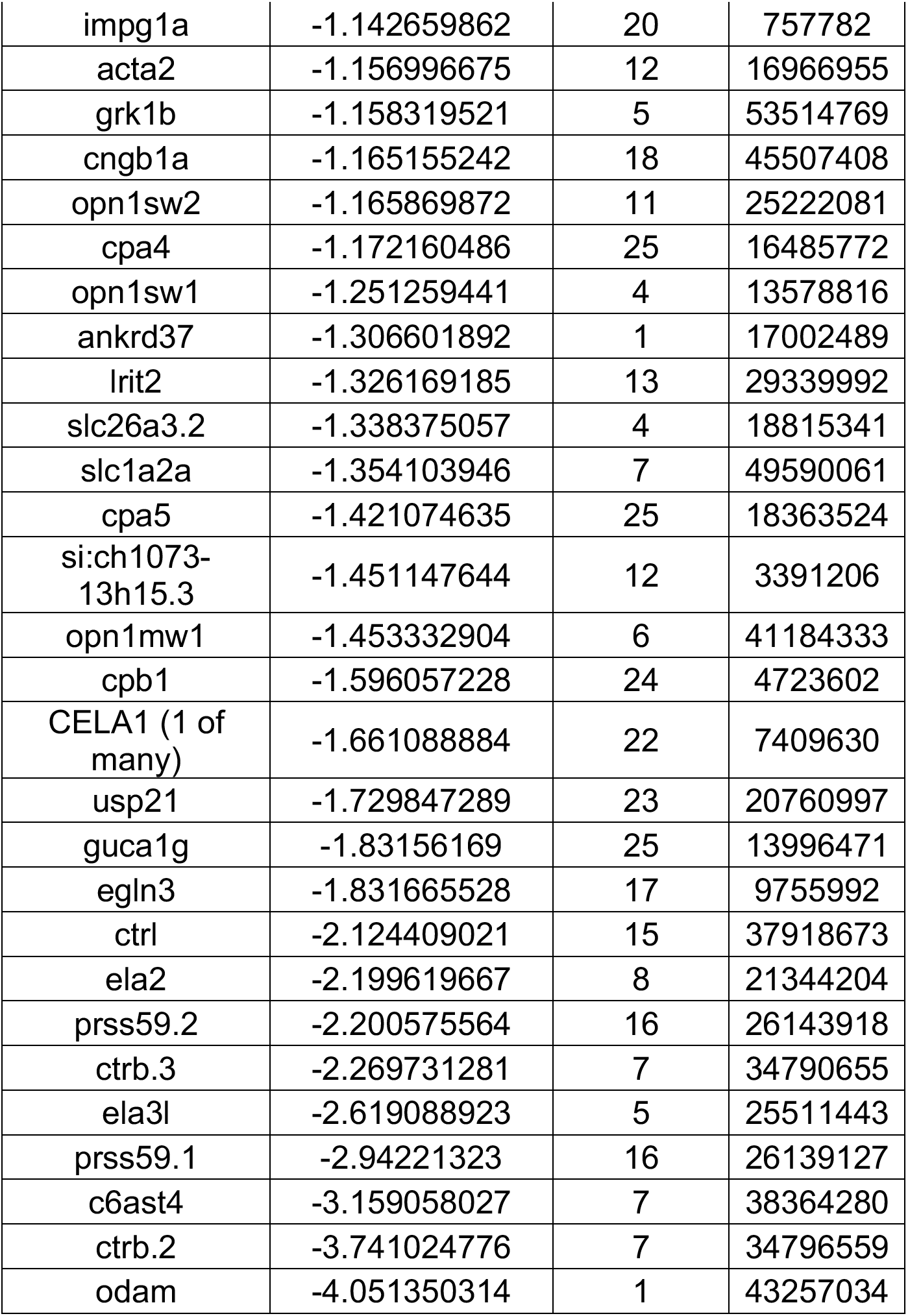
List of transcripts with significant differential expression between wildtype and *yap1^nl13^*mutants.

**Table S2:**
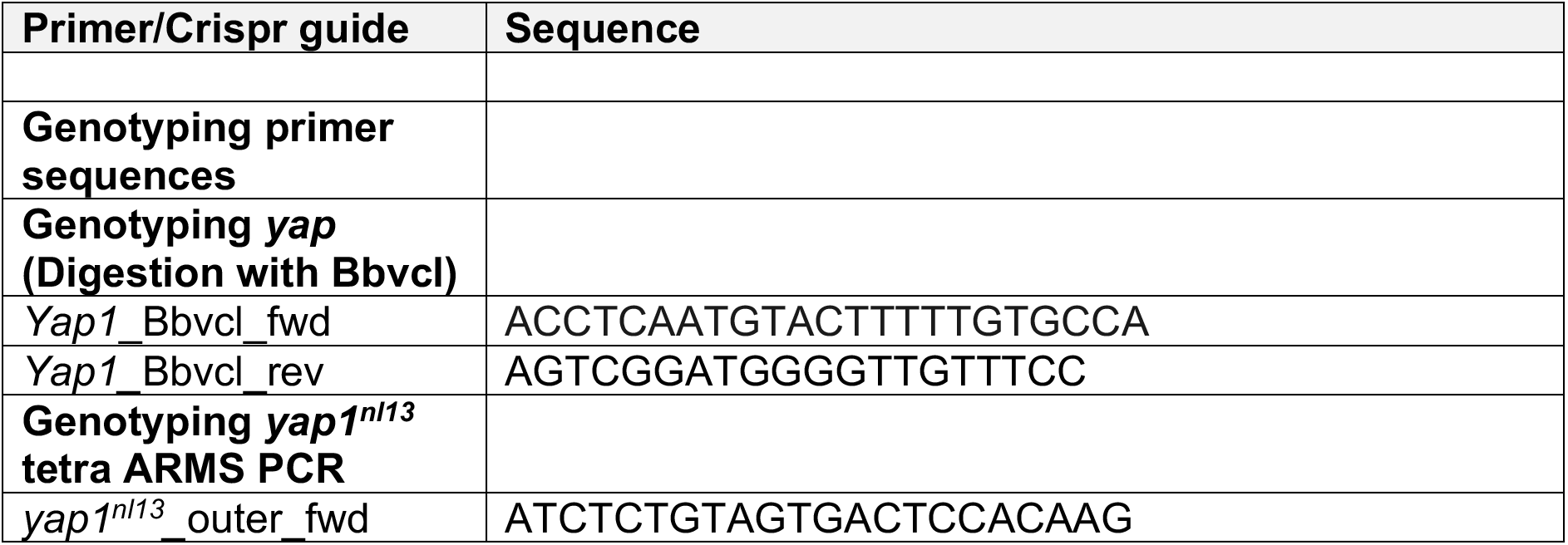
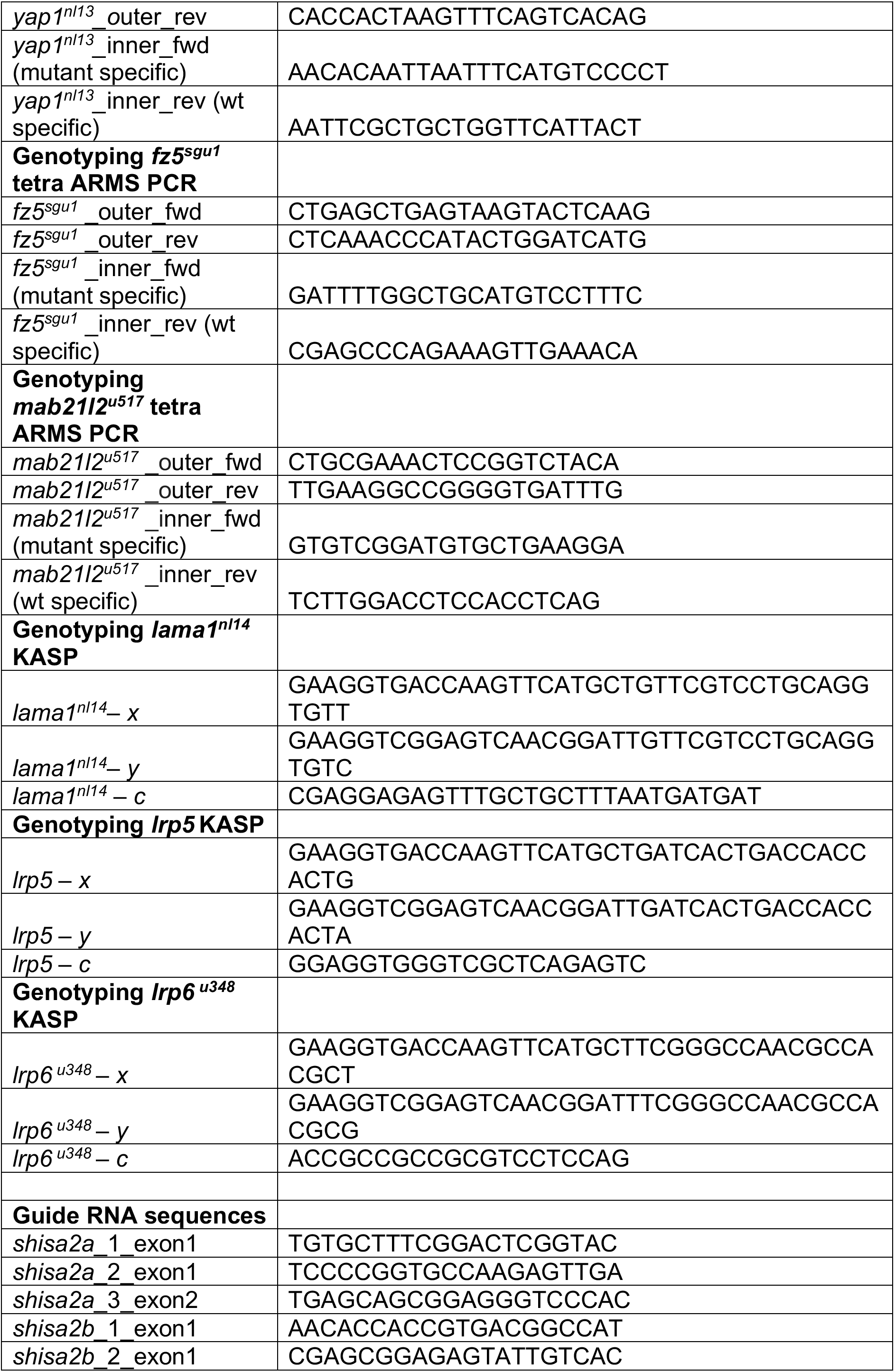
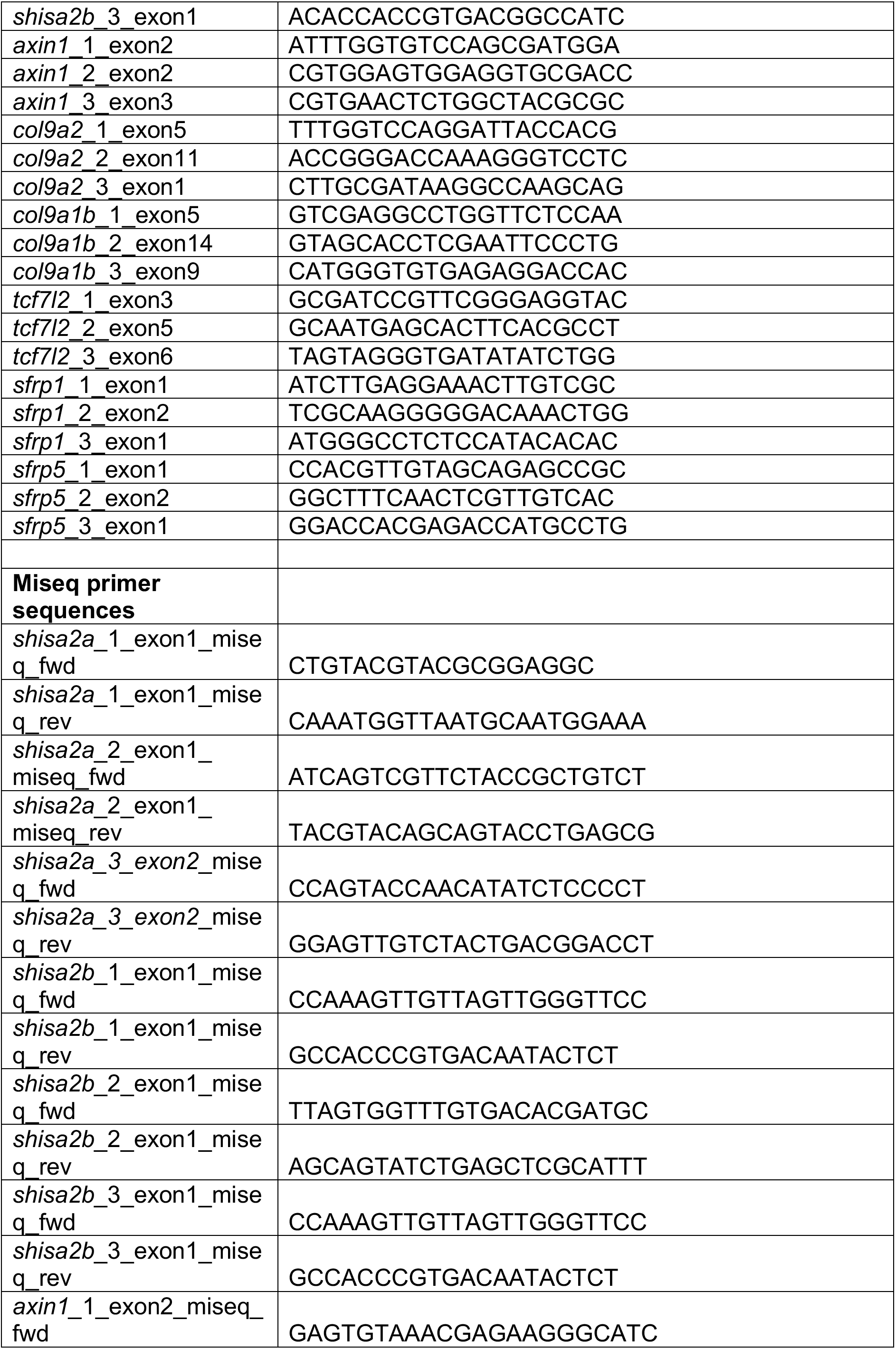
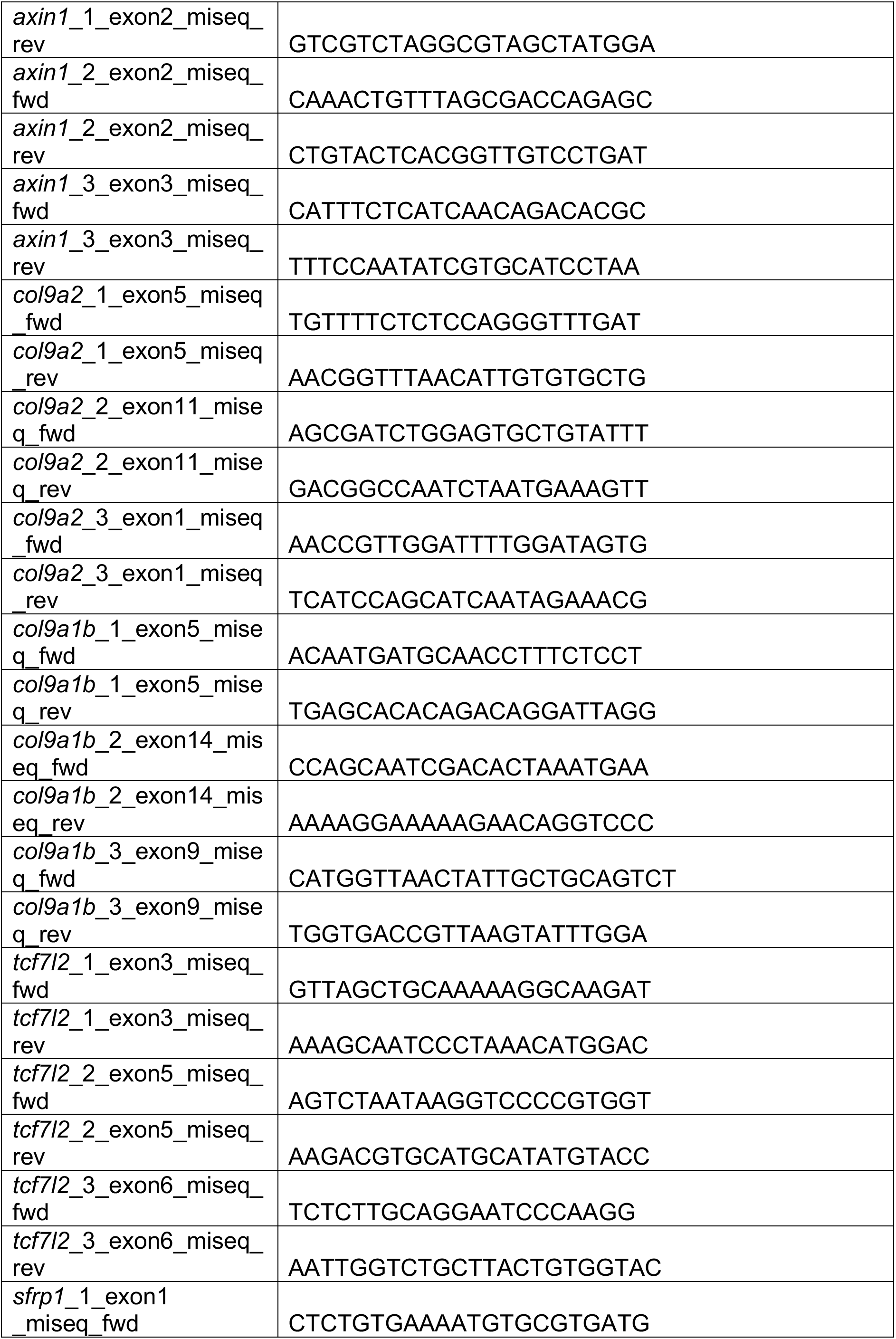
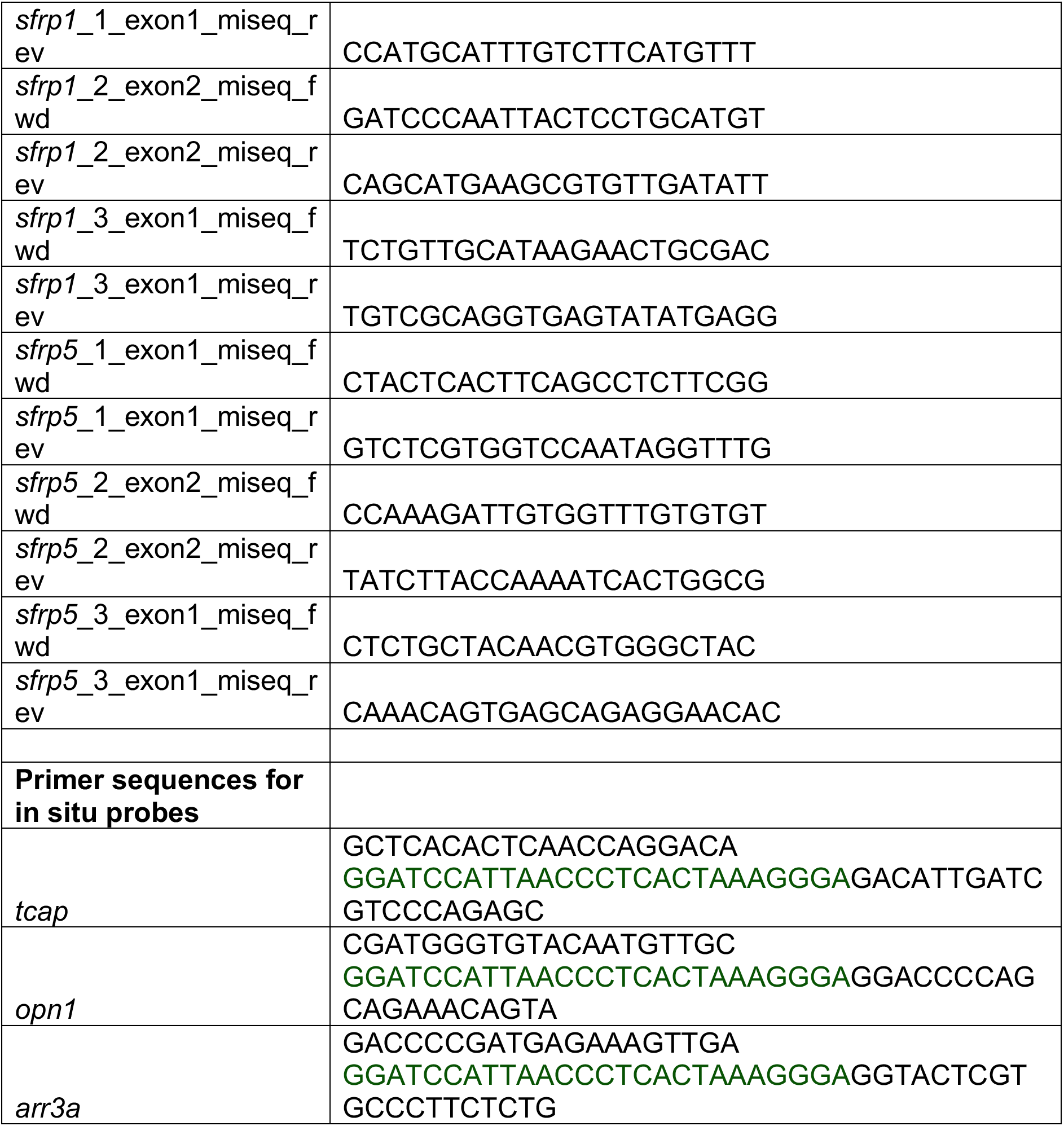
List of primers, gRNAs and probe sequences used in this study.

